# Locus specific epigenetic modalities of random allelic expression imbalance

**DOI:** 10.1101/2021.03.04.433808

**Authors:** Lucile Marion-Poll, Benjamin Forêt, Dina Zielinski, Florian Massip, Mikael Attia, Ava C Carter, Laurène Syx, Howard Y Chang, Anne-Valerie Gendrel, Edith Heard

**Affiliations:** Institut Curie, PSL Research University, CNRS UMR3215, INSERM U934, Paris, France; Directors’ research, European Molecular Biology Laboratory (EMBL), Heidelberg, Germany; Center for Personal Dynamic Regulomes, Stanford University, Stanford, California, USA; Collège de France, Paris, France; Institut Curie, PSL Research University, INSERM U900, Mines ParisTech, Paris, France; Berlin Institute for Medical Systems Biology, Max Delbrück Center for Molecular Medicine, Berlin, Germany

## Abstract

Most autosomal genes are thought to be expressed from both alleles, with some notable exceptions, including imprinted genes and genes showing random monoallelic expression (RME). The extent and nature of RME has been the subject of debate. Here we investigate the expression of several candidate RME genes in F1 hybrid mouse cells before and after differentiation, to define how they become persistently, monoallelically expressed. Clonal monoallelic expression was not observed in ESCs, but when we assessed expression status in more than 200 clones of neuronal progenitor cells, we observed high frequencies of monoallelism. We uncovered unforeseen modes of allelic expression that appear to be gene-specific and epigenetically regulated. This non-canonical allelic regulation has important implications for development and disease, including autosomal dominant disorders and opens up novel therapeutic perspectives.

## INTRODUCTION

Genes are generally thought to be expressed from both alleles, even if their expression is not necessarily equal nor synchronised. There are some exceptions where gene expression occurs exclusively from one allele. This can be due to heterozygous loss-of-function mutations, allelic exclusion via DNA rearrangements (e.g. Immunoglobulins^1^), or epigenetically based differences leading to clonally inherited monoallelic expression. Epigenetically based monoallelic expression is found at parentally imprinted loci^2^, on one X chromosome in females (X-chromosome inactivation)^3^ and for olfactory receptor genes^4^. A less well-understood category concerns autosomal, clonally heritable “random monoallelically expressed” (RME) genes (estimated to comprise 1 to 10% of genes)^5^. RME genes were identified through numerous genome-wide allele-specific analyses of gene expression in single-cell derived clones from hybrid cell lines^6–14^. These studies showed that some genes can be expressed monoallelically from the maternal or paternal allele, or biallelically, and that these expression patterns are inherited during cell division. These RME genes^15^ belong to a wide range of gene ontologies and are enriched in cell surface receptors, developmental regulators, and are often associated with human autosomal dominant diseases^6, 10, 11^.

RME has potentially important implications in development and disease. It might confer many advantages to the organism, such as generating cellular diversity, enhancing adaptability, or regulating gene dosage, but this remains to be investigated^16^. On the other hand, RME may also be detrimental, particularly in the context of heterozygous mutations, as a proportion of cells could express the mutated allele only, thereby contributing to pathological phenotypes in a non-classical manner. Indeed some autosomal dominant disorders involving putative RME loci may be a result of functional nullisomy in a proportion of cells rather than haploinsufficiency^5^.

However, the extent to which clonal and stable RME occurs is still somewhat debated^17, 18^. First, stable RME is often confounded with transient RME observed in single cells, which originates from transcriptional bursting. Although there have been some efforts to identify RME *in vivo*^11, 19^, transient and stable RME cannot be readily distinguished at the single-cell level (by RNA-seq or nascent RNA-FISH) without information about clonality. Second, RME genes are often associated with low expression, raising several issues. A poor detection of lowly expressed genes could lead to unreliability of the allelic measurements. Also, monoallelic expression of a virtually silent gene could arise through incomplete silencing of one allele and subsequent clonal propagation, which would not be functionally relevant. There are therefore critical parameters to take into account, and careful analytical and statistical methods are necessary to confidently identify stable RME, particularly *in vivo*. In addition, the mechanisms underlying stable RME are largely unexplored. In previous studies^6–14^, the number of single-cell derived clones analysed in parallel was rather limited (4-16), rendering any assumptions on the extent of RME and on the mechanisms involved difficult^20^.

Here we assess whether and how some autosomal genes that we previously identified as being RME are prone to persistent and stable monoallelic expression in clones of neural progenitor cells (NPC). We also explore the mechanisms underlying RME and its consequences on gene dosage.

The frequencies of monoallelic expression for a subset of genes involved in disease were assessed in more than 200 newly derived NPC clones. Unexpectedly, we found that genes previously identified as RME show very specific modalities of allelic expression. We propose that these loci are subject to Random Allelic Expression Imbalance (RAExI), with one or several stable states of allelic expression per gene. RAExI consists of a range of different expression modalities, and includes RME as one subtype of RAExI. We also evaluate whether these modalities are determined prior to differentiation. Importantly we assess the consequences of allele-specific expression on gene dosage and on cellular diversity. Finally, we investigate the epigenetic mechanisms involved in the maintenance of differential allelic expression. Taken together, our results reveal unforeseen modes of allelic expression, that appear to be gene specific and epigenetically regulated. This non-canonical allelic regulation affects the genes of this study, which are involved in development and associated with diseases, but the scope of RAExI could be much broader, with large physiological and pathological implications.

## RESULTS

### The study of RME reveals the existence of RAExI

In order to investigate the extent and nature of RME in depth, we selected twelve genes that were previously found to be RME in NPCs (*A2m*, *Acyp2*, *Bag3*, *Cnrip1*, *Eya1*, *Eya2*, *Eya4*, *Grik2*, *Kcnq2*, *Ptk2b*, *Snca*)^11^ or in B-lymphoblastoid clones (*App*)^6^. We also investigated a previously described biallelically expressed gene (*Eya3*). Characteristics of these genes and their association with diseases are summarized in Table 1.

**Table 1.**
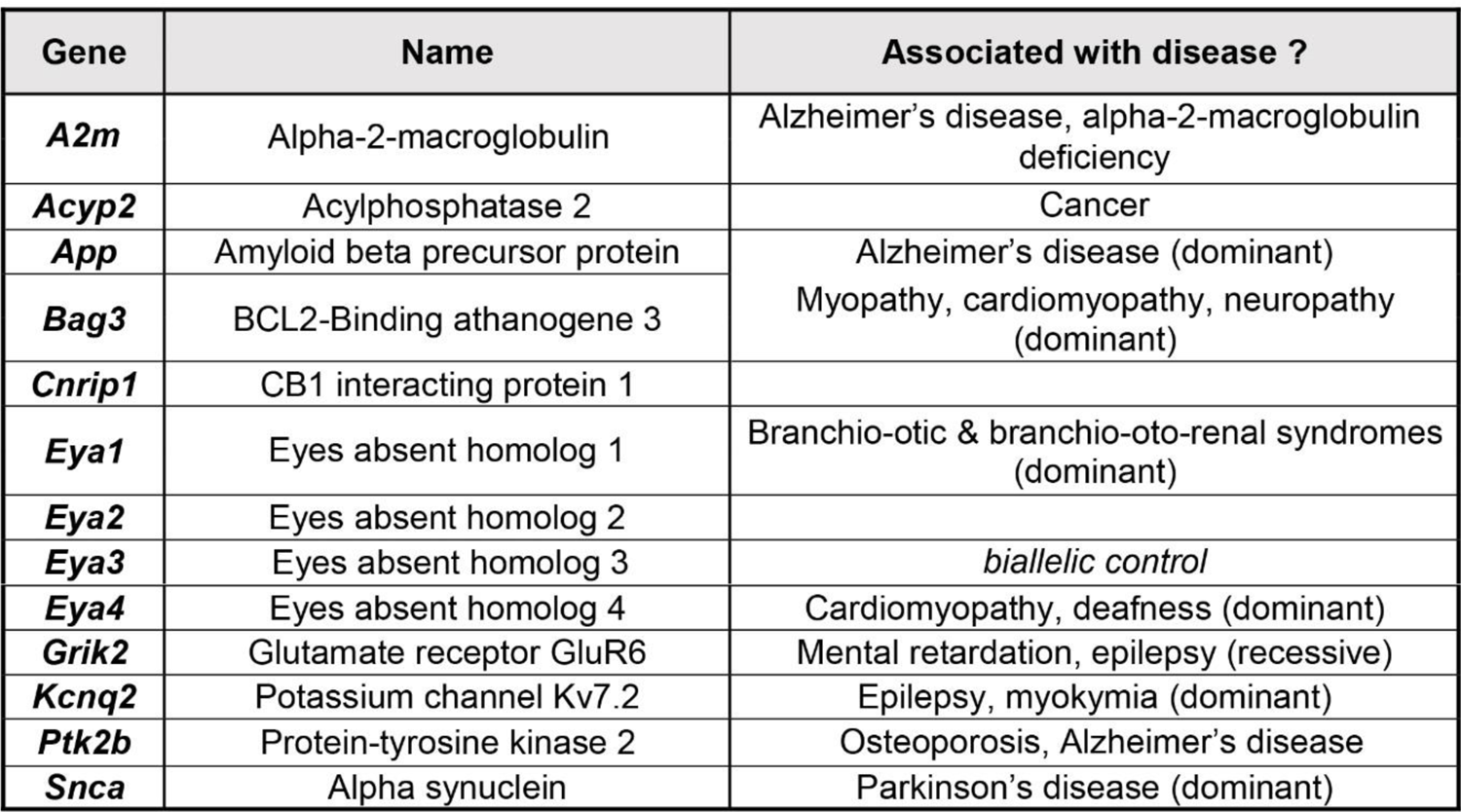
Genes of interest, selected from previous genome-wide studies.

In order to assess the frequency with which these genes show RME, we differentiated male or female hybrid (129/Sv x Castaneus) mouse ES cells (ESCs) into NPCs, as previously described^11, 21^ (Fig. 1, Supplementary Fig. 1). NPCs have the advantage of self-renewing continuously and are clonogenic, as well as being able to differentiate further into neurons, oligodendrocytes or astrocytes *in vitro*^21^. We established 249 independent NPC clonal cell lines from single cells, from six independent differentiation experiments (Supplementary Table 1). The analysis of such a large number of clones confers better statistical power and robustness compared to all previous studies^6–14^.

**Figure 1.**
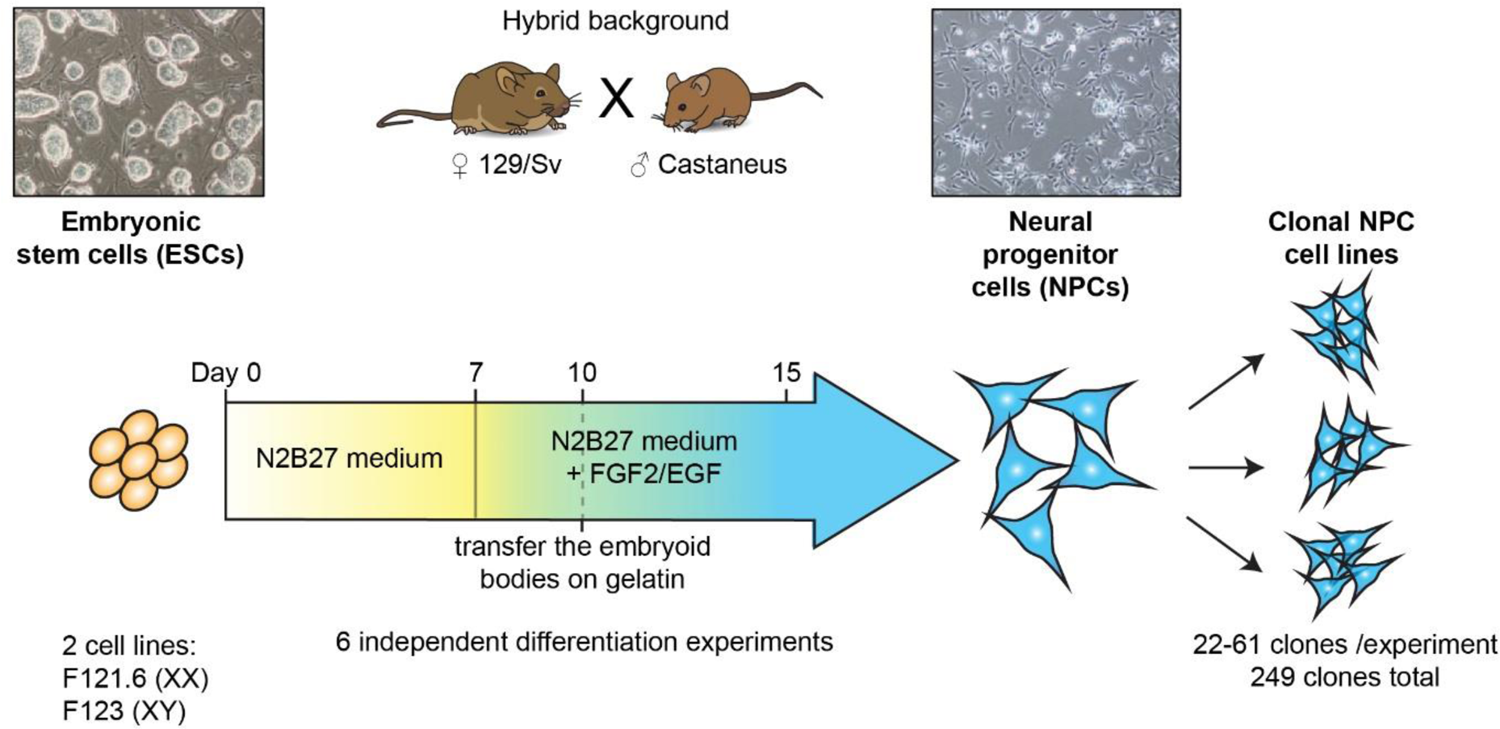
Experimental strategy to assess the frequency of monoallelic expression. Generation of NPC clonal cell lines (249 clones in total) from 2 hybrid ESC lines after 6 independent *in vitro* differentiation experiments.

To evaluate the frequency of monoallelic expression for the candidate genes, we measured their allelic ratios of expression by RT-PCR followed by pyrosequencing in the newly derived NPC clones (65-245 clones/gene, Supplementary Table 2). The allelic expression ratio is defined as the proportion of mRNA expression from the Castaneus (Cast) allele over the total expression. The large number of clones analysed allowed assessment of the distribution of allelic expression ratios across clones for the 13 genes. Strikingly, we observed that the genes have very different profiles (Fig. 2). In order to decompose the distribution of the allelic expression ratio for each gene, we computed Gaussian mixture models to fit the data (see Methods), as this approach can reveal subpopulations of clones and putative allelic expression states. The number of populations for each gene was estimated using the Bayesian Information Criterion (BIC) (Supplementary Fig. 2). We used bootstrapping (see Methods) to assess both the confidence of the number of populations (Supplementary Table 3) and of each Gaussian mixture model component (Supplementary Table 4). This analysis shows that our sample size grants us sufficient power to detect distinct populations. For each gene, we identified one or several subpopulations, defined by a specific mean and variance of the allelic expression ratio (Fig. 2, Supplementary Table 4). Overall, genes display their own modalities of allelic expression.

**Figure 2.**
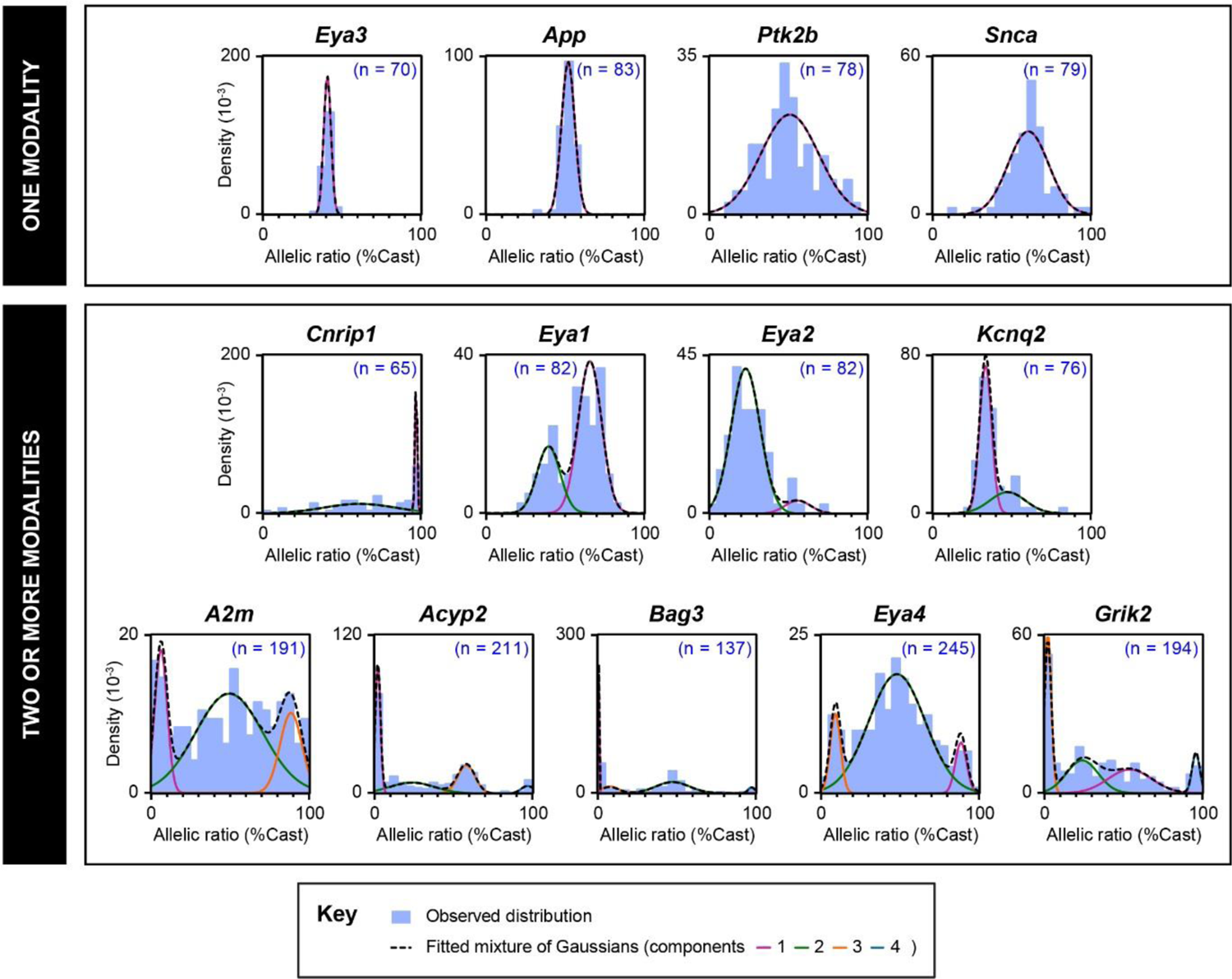
Gene-specific modalities of allelic expression. The distributions of allelic expression ratios in NPC clones are represented as density histograms for the 13 genes of interest. n is the number of NPC clones analysed for each gene. The observed distributions are modelled with Gaussian mixtures. The genes were classified according to the number of subpopulations, computed using the BIC criterion (Supplementary Fig.2).

For *Eya3*, *App*, *Ptk2b* or *Snca,* we found a single population of clones (Fig. 2-top panels), however with very different variances. The biallelic control *Eya3* displays a narrow variance, and a mean allelic ratio of 41% (% Cast) (indicative of a slight genetic bias towards expression of the 129 allele). *App* is biallelic in all tested NPC clones. *Snca* and *Ptk2b* both show a large variance, with the presence of clones showing extreme allelic expression ratios.

For *Cnrip1, Eya1*, *Eya2* and *Kcnq2*, at least two subpopulations could be identified (Supplementary Figure 2, Supplementary Table 3), showing that these genes have more than one modality of allelic expression, each of them with a different variance and degree of allelic expression imbalance (Fig. 2-middle panels). For example, *Cnrip1* is frequently monoallelically expressed from the Cast allele (30% of clones) (Supplementary Table 4), whereas very rare clones display monoallelic expression from the 129 allele (Supplementary Table 2), suggestive of an additional strong genetic bias. *Eya1* on the other hand does not fully exhibit monoallelic expression in any of the clones analysed, but displays two clear subpopulations, biased towards either the 129 or the Cast allele. This observation is unexpected: some genes (such as *Eya1* or *Kcnq2)* do not necessarily show any monoallelic population, but they do show two modalities of expression with different degrees of allelic expression imbalance. They are neither biallelic nor RME genes, but rather display Random Allelic Expression Imbalance (RAExI).

The last group of genes, which includes *A2m*, *Acyp2*, *Bag3*, *Eya4* and *Grik2*, was analysed using additional clones from differentiation experiments 4 to 6, as they appeared to have more complex distributions (Fig. 2, Supplementary Table 2). These genes show three or more modalities of expression (Fig. 2-bottom panels, Supplementary Fig. 2), with gene-specific subpopulations (Supplementary Table 4). They display distinct monoallelic populations in both directions, comprising 18-44% of the clones analysed (Supplementary Table 4), as well as biallelic or biased populations. Thus, *A2m*, *Acyp2*, *Bag3*, *Eya4* and *Grik2* can be considered as RME, RME being a subtype of RAExI. The random monoallelic expression of these five genes is of particular interest as they have been associated with various diseases: *Acyp2* with cancer^22^, *A2m* with Alzheimer’s disease^23^, *Bag3* with myopathy^24^, *Eya4* with deafness^25^ and *Grik2* with epilepsy^26^. *Acyp2* and *Bag3* show the highest proportion of monoallelically-expressing NPC clones, with small variance, and thus represent good candidates to further explore the features of allele-specific expression.

### RAExI is established and maintained during differentiation

To determine whether the different states of allelic expression observed in NPCs arise during differentiation or are already present in ESCs, we established 18 ESC clones from the hybrid female ESC line and measured allele-specific expression of *Acyp2* and *Bag3* using RT-PCR and pyrosequencing. Both genes were found to be biallelically expressed in all ESC clones tested (Fig. 3a, left panels), in contrast to NPCs, as shown using cumulative distributions of the allelic expression ratios in both cell types (Fig. 3a, right panels). This indicates that allele-specific expression is established during differentiation.

**Figure 3.**
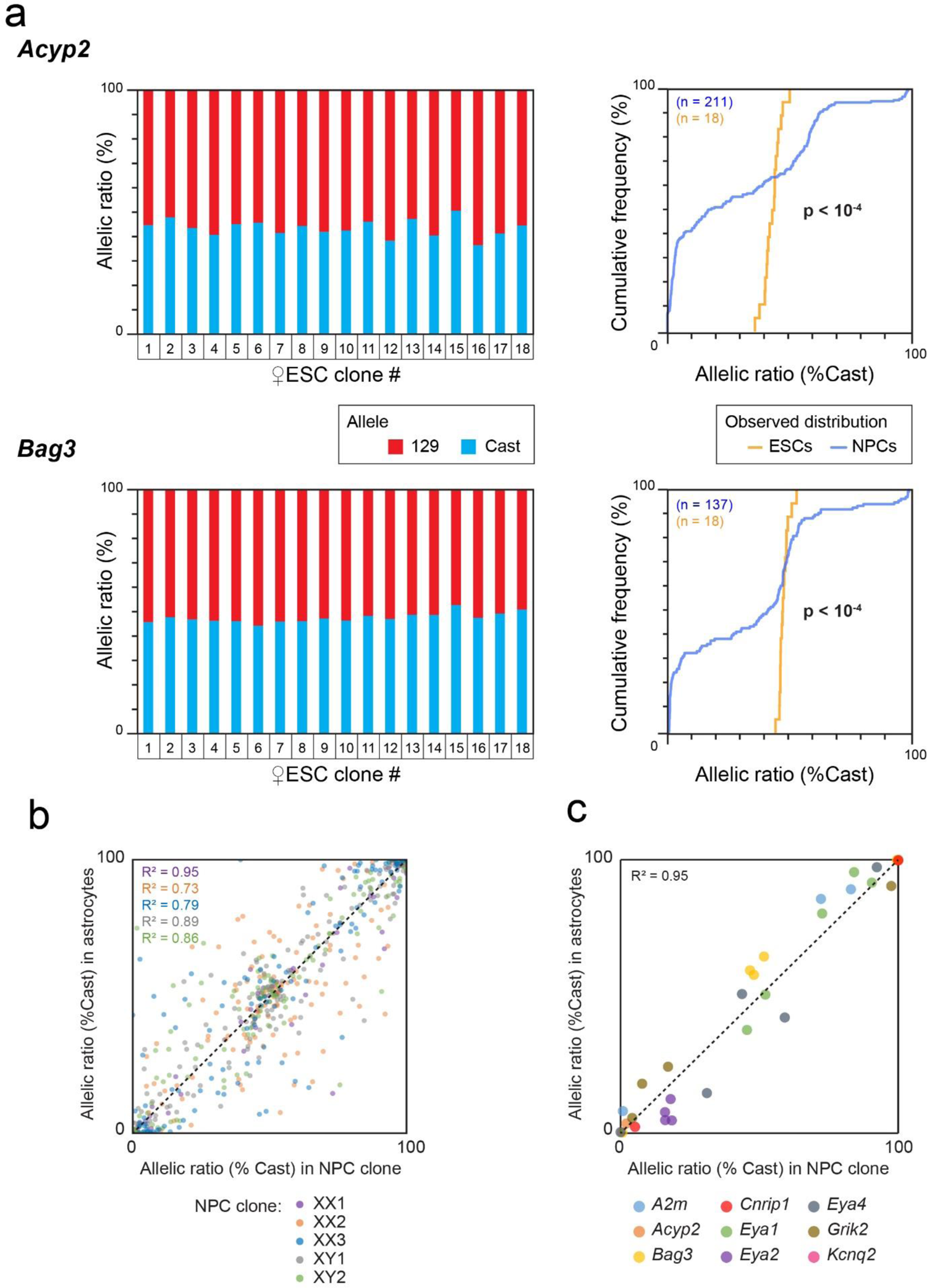
Allele-specific expression is established during differentiation and then maintained. (a) Analysis by RT-PCR followed by pyrosequencing of *Acyp2* and *Bag3* allelic expression ratio in 18 clones generated from the F1-21.6 ESC line (left panels). Comparison of the cumulative distributions of allelic expression ratios in ESCs and NPCs (Kolmogorov-Smirnov tests, right panels). (b) Comparison of the allelic expression ratios of the genes previously categorised as RME in NPCs^11^ and after differentiation into astrocytes, for 5 NPC clones, measured by allele-specific RNA-seq. The number of assessable genes per clone (values with less than 20 reads were excluded) is: n_XX1_=75, n_XX2_=146, n_XX3_=121, n_XY1_=150, n_XY2_=137. Pearson correlation coefficients were calculated for each NPC clone. (c) Comparison of the allelic expression ratio for the 9 genes, which showed several modalities of expression, in 5 NPC clones and after differentiation into astrocytes.

We also evaluated the stability of allele-specific expression following further differentiation of NPCs to astrocytes. We previously showed that the RME pattern of 6 genes is maintained over cell passaging, and following differentiation of two independent NPC clones to astrocytes^11^. Here, we expand this analysis by performing allele-specific RNA-seq of 5 NPC clones after differentiation into astrocytes (Supplementary Fig. 3). We observed that the allelic expression ratios for the genes previously listed as RME in NPCs^11^ are globally stable following differentiation (Fig. 3b). Likewise, the allelic expression ratios of our RAExI genes appear stable (Fig. 3c). Overall, these analyses confirm and expand previous observations^10, 11^ that allelic expression imbalance is established during differentiation of ESCs to NPCs and stably maintained once established.

### RAExI gives rise to stochastic diversity

We next explored whether the allelic choices established during differentiation of ESCs to NPCs are in any way coordinated for multiple independent genes. This might occur if a common mechanism were involved in establishing or maintaining allele-specific expression of different genes, in particular for genes belonging to the same family (such as the *Eya* gene family). In order to reveal any potential concomitance in the allelic choices of the different genes, we performed hierarchical clustering of the allelic expression ratios for our 13 selected genes in the 44 NPC clones for which we had pyrosequencing data for all genes (Fig. 4a). We found that each NPC clone has a unique combination of allelic expression ratios for different genes and that none of the genes were coordinated, indicating that genes make allelic decisions independently of each other.

**Figure 4.**
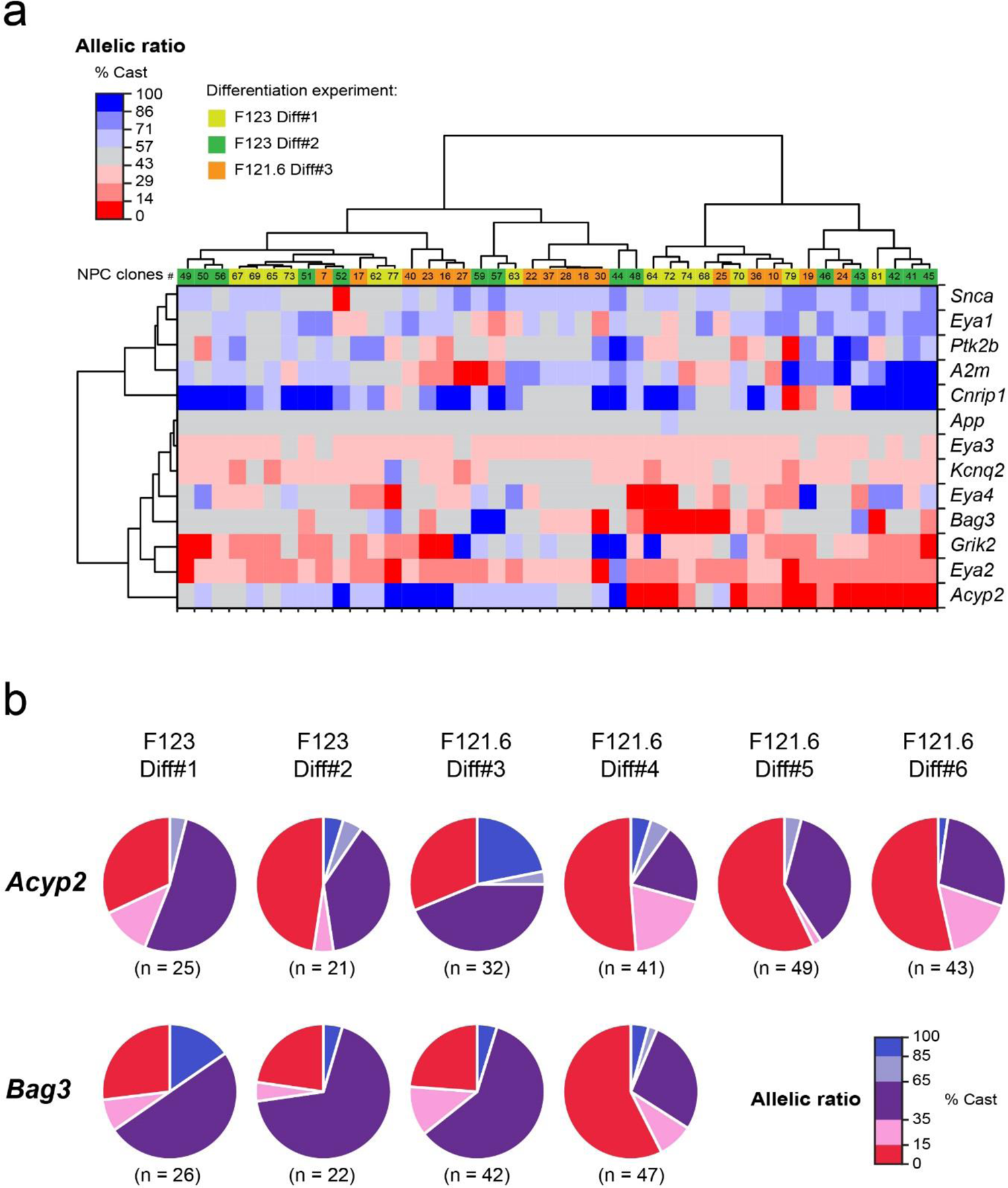
Allelic imbalance generates cellular diversity. (a) Hierarchical clustering heatmap of allelic expression ratios for 13 genes in 44 NPC clones. (b) Proportions of NPC clones in each category of allelic expression status for *Acyp2* and *Bag3* for 6 independent differentiation experiments. n is the number of clones analysed for each gene and each differentiation.

The NPC clones in this study were generated following six independent differentiation experiments of two ES cell lines. In order to assess whether the allelic choice for a given gene is deterministic, we compared the outcomes of all experiments. We observed that the allelic choices can be highly variable between differentiation experiments, notably for *A2m*, *Acyp2, Bag3*, *Cnrip1*, *Eya2,* and *Eya4* (Fig. 4b, Supplementary Fig. 4). These observations indicate that the allelic choice is more consistent with a stochastic establishment during differentiation. The multiple combinations of allelic expression patterns that appear during differentiation therefore give rise to an unpredictable high level of cellular diversity.

### RME is associated with reduced gene expression dosage

We next explored the link between monoallelic expression and gene dosage by measuring the expression levels by RT-qPCR for the RME genes *Acyp2*, *Bag3* and the biallelic gene *Eya3* in all NPC clones. While the biallelic control *Eya3* shows similar expression levels in all clones, *Acyp2* and *Bag3* could be divided in two main groups, showing either high or low expression levels (Fig. 5a). Interestingly, we found that there are less monoallelically expressing clones in those with low expression than in the group of clones showing high expression, indicating that monoallelic expression is not necessarily associated with low expression. Moreover, in clones with high expression levels, we found that the quantity of mRNA is reduced by half in monoallelic compared to biallelic clones. This impact on dosage is also observed at the protein level for *Bag3*, as we found a highly significant linear correlation between mRNA and protein levels (Fig. 5b, Supplementary Fig. 5). BAG3 is a multifunctional co-chaperone protein that regulates diverse biological processes such as apoptosis and autophagy. It is a dosage-sensitive gene as its overexpression has been shown to promote apoptosis^27^, while its downregulation attenuates it^28^. RME appears to be a way to down-regulate protein levels, and thus could have consequences on cellular functions.

**Figure 5.**
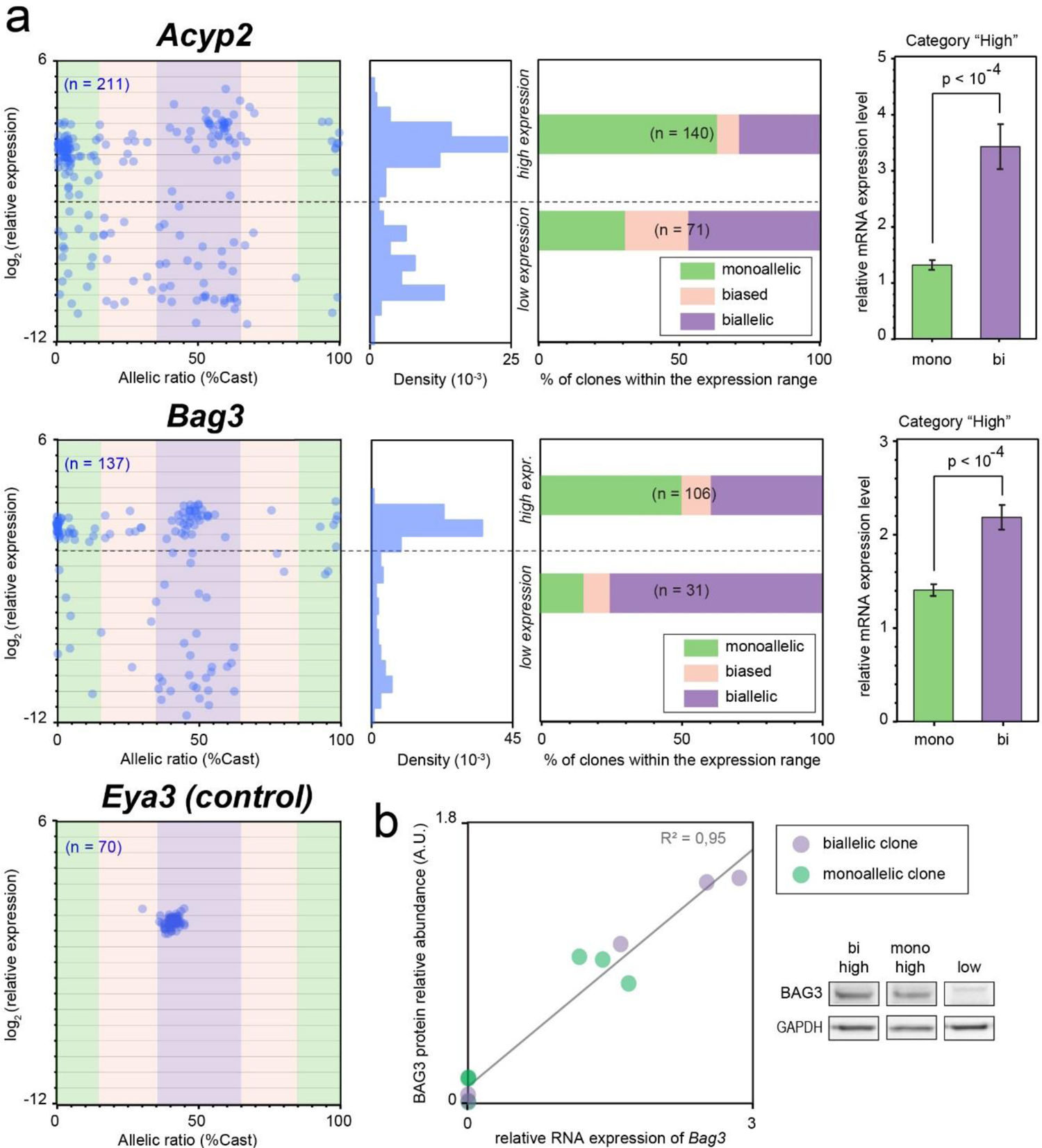
Random monoallelic expression is associated with reduced gene dosage. (a) Comparison of the allelic expression ratio and mRNA expression levels (measured by qPCR), for *Acyp2*, *Bag3* and *Eya3*, in NPC clones (n is the number of clones analysed for each gene; left panels). The density histograms show the distribution of expression levels; the dashed lines indicate the threshold used to categorise high or low expressing clones (middle left panels). Proportion of clones in each allelic category for clones showing high or low expression levels (middle right); allelic categories: monoallelic [ratio<15% or >85%], biased [ratio 15-35% or 65-85%], biallelic [ratio 35-65%]; n is the total number of clones. mRNA levels in monoallelic clones compared to biallelic clones (excluding the clones with low expression levels). Data are means ±SEM; two-tailed unpaired t-tests, *Acyp2* t=6.5 df=131, *Bag3* t=5.6 df=92 (right panels). (b) BAG3 relative protein levels measured by western blot (normalised to GAPDH levels), compared to the relative RNA abundance measured by RT-qPCR, for a selection of 11 NPC clones. Linear regression *F*_(1, 9)_ = 176, *p* < 10^-4^. A western blot image of the Bag3 full-length form is shown for 3 representative NPC clones.

### Cell-type specificity of RME

We next addressed the potential impact of RME *in vivo* for some of our candidate genes, at the adult stage. We established a new protocol (see Methods), which allows the detection of nascent RNA in the adult brain, known for accumulating age-related autofluorescent molecules and lipofuscin-like pigments. I*n vivo* analyses are generally limited by problems arising from cellular heterogeneity, transcriptional bursts, and detection efficiency. We therefore applied a statistical test to evaluate whether or not we could rule out the possibility that the number of monoallelic cells observed by RNA-FISH would simply be a consequence of transcriptional bursts or a lack of detection (see Methods). To perform this analysis, both alleles must be genetically identical (inbred background), and the cells analysed must be homogeneous (i.e. same cell type, with the same kinetics of transcriptional bursts and the same probability to detect an expressed allele for a given gene).

In the brain, different cell types are usually intermingled, except in some specific regions, such as the hippocampus, where the somas of homogeneous cell types are tightly packed, making it an ideal model to investigate RME *in vivo*. Among the genes studied here, *App* and *Grik2* are specifically relevant to study in this region, as they are respectively associated with Alzheimer’s disease and epilepsy, two pathologies that affect the hippocampus. We performed nascent RNA-FISH for *App* and *Grik2* on hippocampal sections of adult inbred (C57Bl/6J) mice, and determined the percentage of cells showing monoallelic expression in the pyramidal neurons of the CA1 and CA3 regions. We observed that *Grik2* is detected as monoallelic in 30% of cells on average in CA1, unlike in the neighboring CA3 region where *Grik2* is biallelic, whereas *App* is biallelically expressed in both CA1 and CA3 neurons (Fig. 6). The percentage of CA1 neurons expressing *Grik2* monoallelically is similar for the 3 animals analysed, and the observed distribution is significantly different from what would be expected from a biallelic gene showing bursty expression or with incomplete detection (see Methods). This indicates the presence of stable monoallelic expression *in vivo*. These results parallel the situation in NPCs, where *App* is biallelic and *Grik2* shows RME (with 39% of the clones showing monoallelic expression) (Supplementary Table 4). Even though it was found to be monoallelic in lymphoblastoid cell lines^6^, *App* is not monoallelic in our study, indicating that this dosage-sensitive gene is not regulated in an allele-specific manner in hippocampal mouse neurons or NPCs. Monoallelic expression of *Grik2* appears to be cell-type specific in the hippocampus *in vivo*. Interestingly, these findings are consistent with the observation that *Grik2* tends to be more highly expressed in CA3^29^. Moreover, it has been shown that increased expression in CA1 pyramidal cells is deleterious as it can trigger seizures^30^. Thus, RME may contribute to the regulation of gene dosage both *in vitro* and potentially *in vivo*.

**Figure 6.**
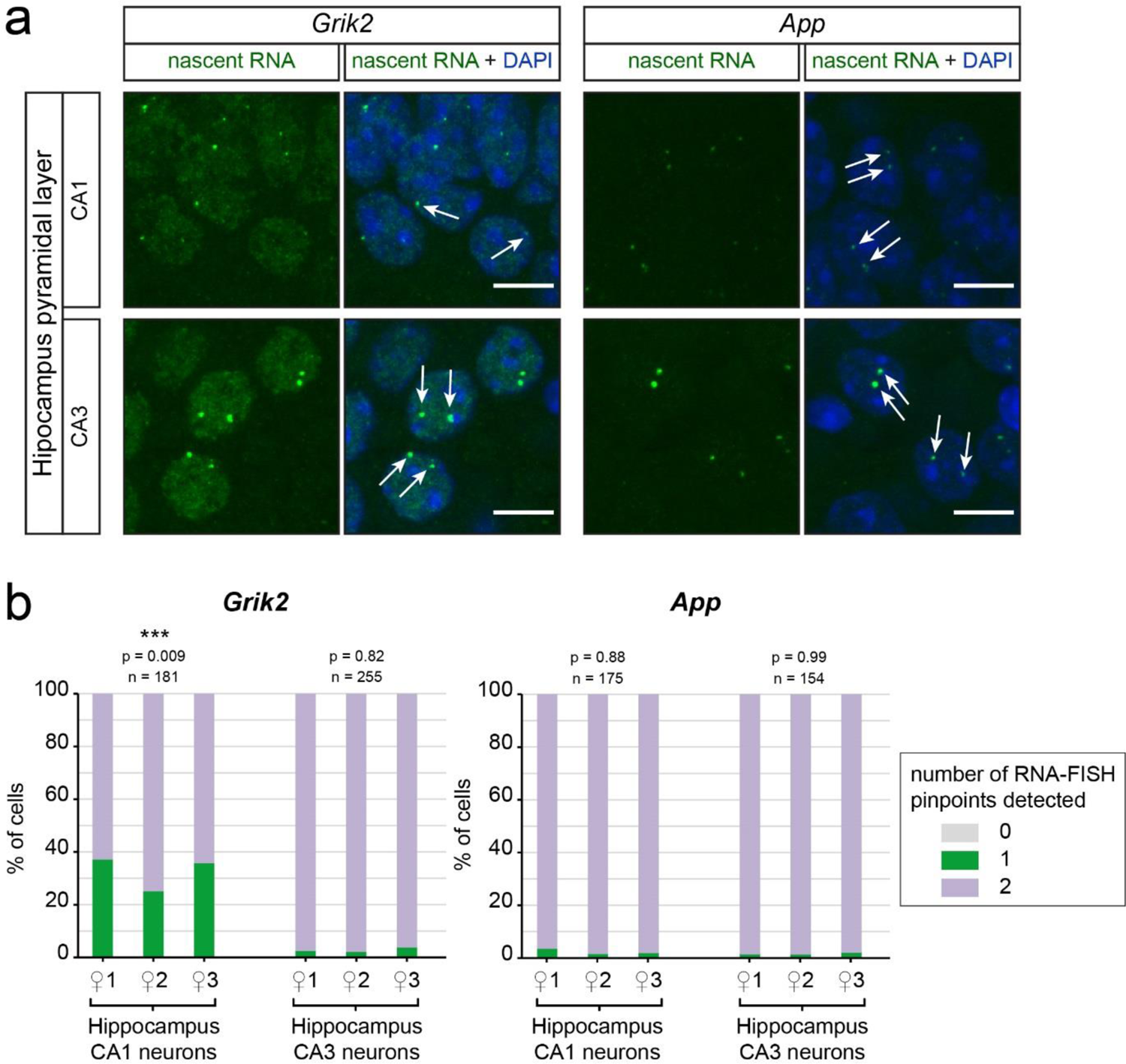
Monoallelic expression is cell-type specific *in vivo*. (a) Z-stack of confocal images showing representative cells with nascent RNA-FISH signals for *Grik2* and *App*, in hippocampal sections of adult inbred mice. Scale bar 10µm. (b) Quantification of nascent-RNA FISH in pyramidal neurons of CA1 and CA3, in 3 female inbred mice. Chi² test (see Methods); n is the number of cells analysed for each gene and each region.

### Local regulation of RME

To explore how allele-specific expression could be regulated, we examined whether neighbouring genes to loci showing RME (*A2m*, *Acyp2*, *Bag3*, *Eya4*, *Grik2,* Fig. 2) also showed RME in the same clones, or some degree of RAExI. Common trans-acting factors could control several genes sharing cis-elements within a genomic region. We used allele-specific RNA-seq datasets from 16 NPC clones (9 previously published and 7 additional clones from this study - Supplementary Table 5)^11, 31^ and compared the distribution of allelic expression ratios for the *Acyp2*, *A2m*, *Bag3*, *Eya4* and *Grik2* genes, together with eight neighbouring genes located upstream and downstream (Fig. 7a, Supplementary Fig. 6a). We found that neighboring genes generally show biallelic expression, with a few exceptions that display uncorrelated allelic imbalance (*Apobec1*, *Slc18b1*). This indicates that the RME is not regulated within a domain but is rather an intrinsic property of each gene, in agreement with previous observations^11, 13^.

**Figure 7.**
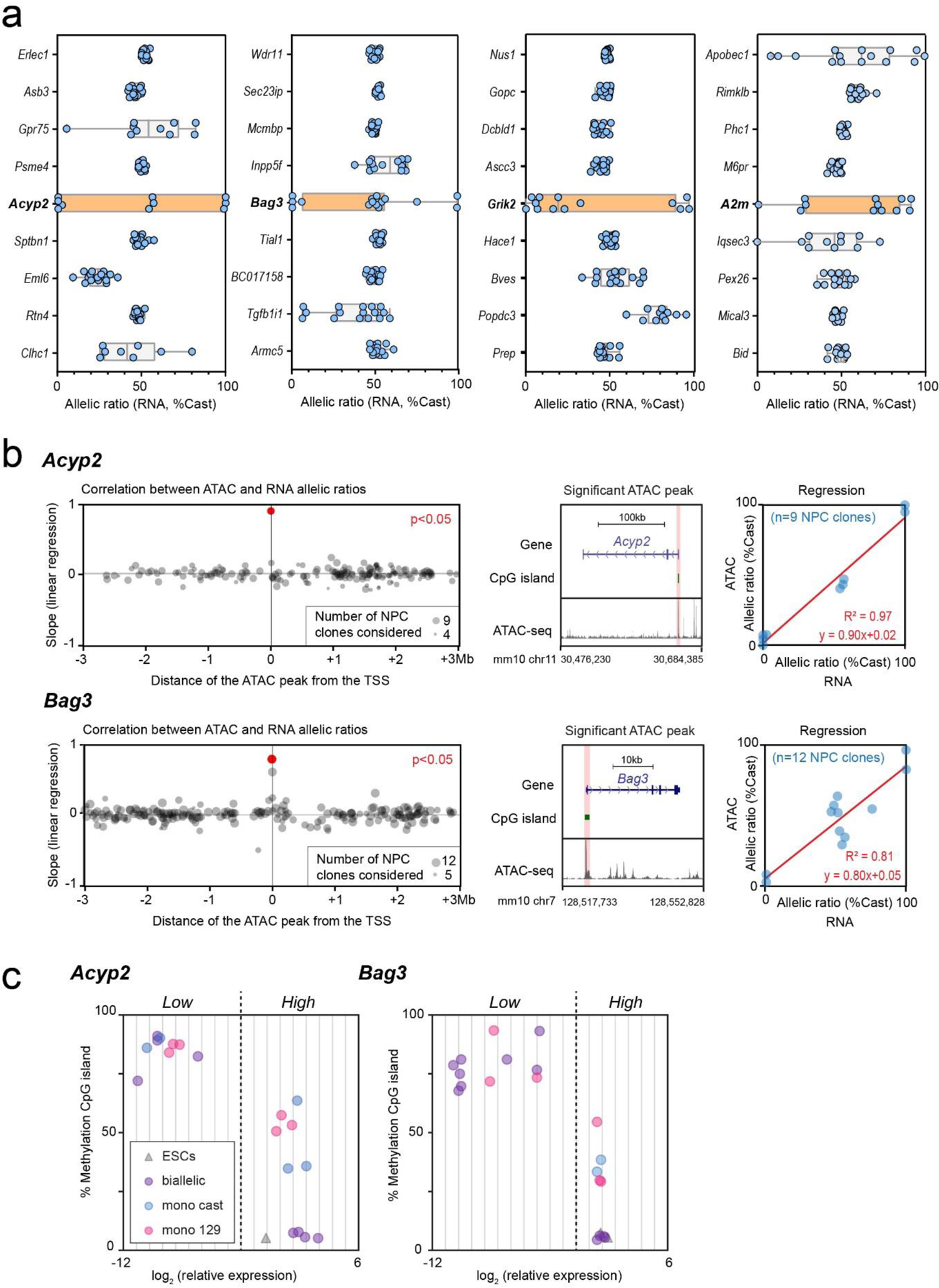
The allelic imbalance is regulated locally. (a) Comparison of the distribution of allelic expression ratios for *Acyp2*, *Bag3*, *Grik2* and *A2m*, with 4 upstream and 4 downstream neighbouring genes, measured by allele-specific RNA-seq in 16 NPC clones. The orange color highlights the whisker plot of the RME gene of interest. (b) Identification of the genomic region where allele-specific accessibility correlates best with expression, by linear regression between the ATAC-seq allelic accessibility ratio and RNA-seq allelic expression ratio, for ATAC-seq peaks (containing at least one SNP) within a region of ±3 megabases around the TSS. *Pvalue* adjusted with Bonferroni correction, number of ATAC peaks: 177/207 *Acyp2*/*Bag3* (left panels). Representative ATAC-seq track along the gene, with highlighting of the ATAC-seq peak, located at the TSS, that correlates significantly with the RNA-seq (middle panels). Linear regression showing the correlation between the ATAC-seq allelic accessibility ratio and the RNA-seq allelic expression ratio at the TSS (right panels). ATAC-seq data and RNA-seq datasets are from 13 NPC clones. *Acyp2 F_(1,7)_* = 227, *p* = 1.3 x 10^-6^; *Bag3* F_(1,9)_ = 42, *p* = 7.3 x 10^-5^. (c) DNA methylation levels over the CpG island located at the TSS of *Acyp2* and *Bag3*, measured by Sequenom bisulfite analysis in 20 representative NPC clones and the 2 ES cell lines, characterised by high or low expression levels and mono- or biallelic expression.

We then aimed at identifying the regulatory regions controlling allelic imbalance expression of the genes validated as RME (*A2m*, *Acyp2*, *Bag3*, *Eya4* and *Grik2*). We used allele-specific ATAC-seq datasets available for 13 NPC clones (Assay for Transposase-Accessible Chromatin)^13^, and RNA-seq for the same clones (7 previously published and 6 newly sequenced - Supplementary Table 5). In a previous study^13^, in 7 NPC clones for which both RNA-seq and ATAC-seq data were available, RME genes were found to be associated with random monoallelically accessible elements located mostly at promoters. However, this study was carried out in a genome-wide manner without investigating our individual RME genes of interest.

We analysed all peaks of accessible chromatin with allele-specific information within a 6 megabase region surrounding each gene (see Methods). For each peak, we determined whether the allele-specific chromatin accessibility correlated with allele-specific expression, by linear regression (Fig. 7b, Supplementary Fig.6b,6c -left panels). Even though not all peaks had allelic information, we found a genomic region that shows significant correlation with allelic expression for *A2m*, *Acyp2 Bag3* and *Grik2*, located in each case at the transcription start site (TSS) (Fig. 7b, Supplementary Fig.6c, middle and right panels). This is indicative of a local regulation of RME, in line with the observation that random monoallelically accessible elements are enriched at promoters^31^.

### Epigenetic maintenance of RME

*Acyp2* and *Bag3* showed the highest correlation at the TSS between chromatin accessibility and expression (Fig. 7b). The allele-specific accessible region at the TSS overlaps with a well-defined CpG island (CGI). We therefore measured DNA methylation levels in 20 representative NPC clones and in ESCs using Sequenom analysis of bisulfite-treated DNA. We found that CGI are hypermethylated in clones where the gene is silenced or lowly expressed, hypomethylated in biallelically-expressing clones and in ESCs, and show an intermediate level of methylation in clones with monoallelic expression (Fig. 7c). This suggests that the alleles that are lowly expressed become methylated during differentiation and that DNA methylation correlates with expression levels, rather than directly with the allelic expression status.

To further investigate the role of DNA methylation, as well as other epigenetic mechanisms in the maintenance of RME patterns in NPCs, we designed a screen using a library composed of 181 drugs targeting epigenetic modifications or pathways (Supplementary Table 6). We applied the screen to *Bag3* as a proof-of-principle, as it is a particularly interesting target for allele-specific regulation, due to its multiple functions, its dosage sensitivity and its implications in disease. First, we measured the impact of these molecules on *Bag3* allelic expression in two selected female NPC clones (#31 and #84, expressing *Bag3* from the 129 allele or the Cast allele, respectively), after two days of treatment at a standard concentration of 10 µM (see Methods). We identified several epidrugs that induce an increase in the ratio of expression of the silent allele in both clones (Fig. 8a). A secondary test was conducted using the 11 hits that showed the strongest shift on both NPC clones, on a higher number of cells from NPC clone #84 (Fig. 8b). This analysis confirmed that treatment with the DNA methyltransferase inhibitor Decitabine, as well as two histone deacetylase (HDAC) inhibitors (Dacinostat and CUDC-101) significantly increase the *Bag3* ratio of expression of the silent allele. These drugs inhibit transcriptional repression mechanisms. In order to refine the concentration, we performed a dose-response of Decitabine and Dacinostat, with fresh compounds. The two drugs showed a dose-dependent effect, and reached a maximal effect at 10 µM (Fig. 8c), with an EC50 of 1.6 µM and 0.9 µM, for Decitabine and Dacinostat respectively. Finally, we confirmed their effect on both clones, with a 10 µM treatment reduced to 24 h (Fig. 8d). These analyses show that the silent allele of *Bag3* can be reactivated, and thus it importantly shows that RME is epigenetic, rather than caused by genetic aberrations or gene rearrangements. The maintenance of monoallelic expression of *Bag3* in NPCs relies on a combination of epigenetic modifications, including DNA methylation and histone deacetylation, which are actively involved in the inheritance of the transcriptional state. The inhibition of such pathways can be used to reexpress the silent allele, offering new perspectives for therapeutic strategies for RME genes associated with autosomal dominant disorders.

**Figure 8.**
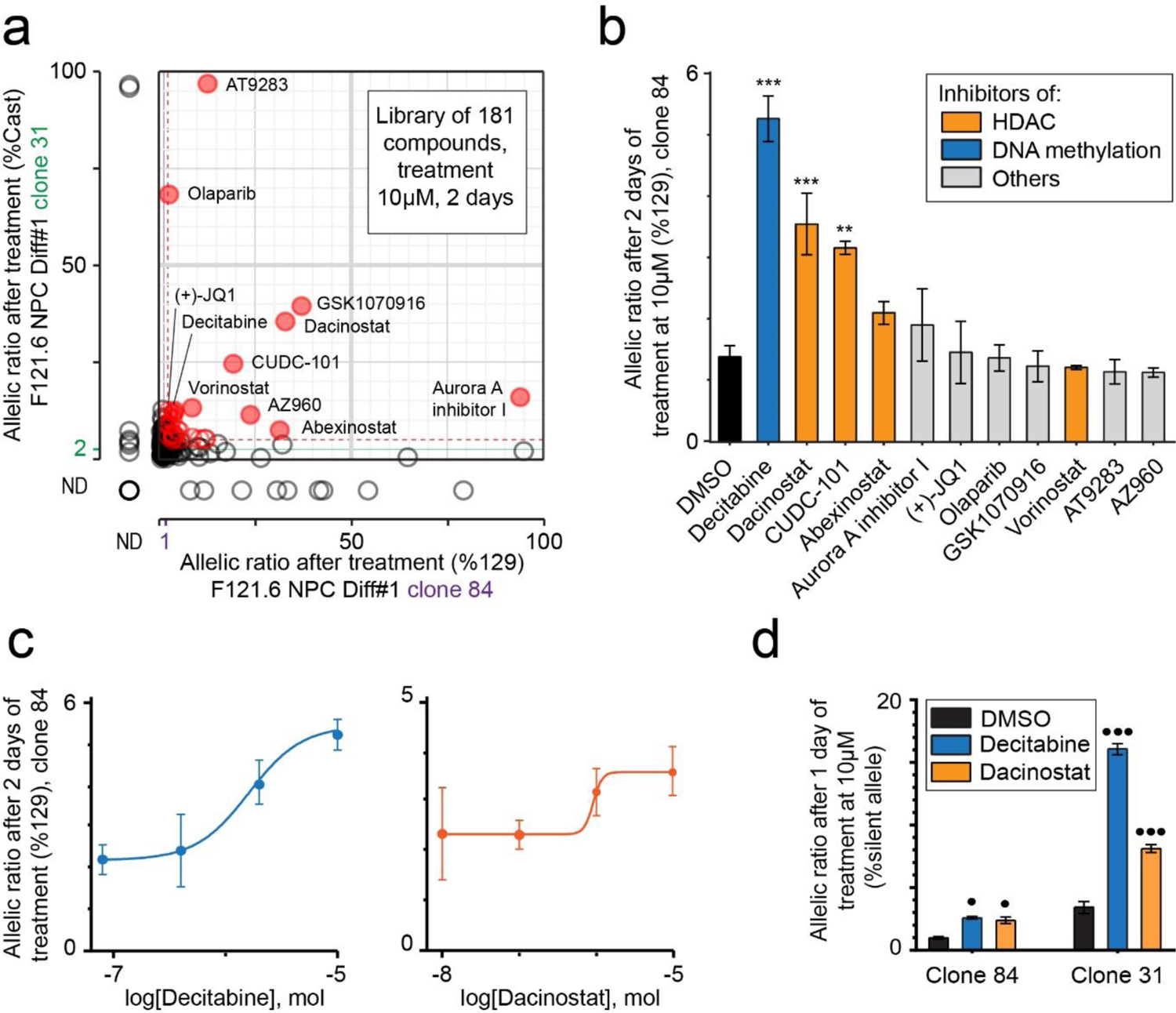
Epigenetic maintenance of the monoallelic expression of *Bag3*. (a) Effect of treatment with an epidrug library (2 days at 10 µM) on the allelic expression ratio of two NPC clones (#31 & #84), measured by pyrosequencing following RT-qPCR. Purple and green lines represent the mean of the 11 controls, dashed red lines are mean + 3 x Stdev, red circles are compounds with an effect > mean + 3 x StDev for both clones. NV, no value (b) Validation of 3 compounds among the 11 best hits from (a) on NPC clone #84 (2 days of treatment at 10 µM). DMSO is used as the vehicle. Data are means ± SEM (*n* = 3 replicates per condition). One-way ANOVA: treatment effect, *F*_(11, 40)_ = 12, *p* < 10^-4^. Dunnett’s *post hoc* test: \**p* < 0.05, \*\*\**p*< 10^-4^ (DMSO versus treatment). (c) Dose-response curve for Decitabine and Dacinostat for NPC clone #84, after 2 days of treatment. Data are means ± SEM. (d) Confirmation of the effect of Decitabine and Dacinostat on NPC clones #31 and #84, after 1 day of treatment at 10 µM. DMSO is used as the vehicle. Data are means ± SEM (*n* = 3 replicates per condition). Two-way ANOVA: treatment effect, *F*_(2, 18)_ = 200, *p* < 10^-4^; clone effect *F*_(1, 18)_ = 548, *p* < 10^-4^; interaction, *F*_(2, 18)_ = 120, *p* < 10^-4^. Bonferroni *post hoc* test: ^•^*p* < 0.05, ^•••^*p* < 10^-4^ (DMSO versus treatment).

## DISCUSSION

The main goal of our study was to explore whether and how some autosomal genes are prone to persistent and stable monoallelic expression, by focusing on a subset of genes of interest. The analysis of a very high number (>200) of NPC clones revealed that genes previously identified as RME actually display one or several modalities of allelic expression, with different degrees of allelic imbalance. These observations unexpectedly uncovered the existence of RAExI. In light of our findings, RME appears to be a specific case of RAExI.

In this study, we used NPCs to assess RAExI and we showed *in vivo* that this phenomenon can be cell-type specific. It will be important to expand the study of RAExI to different cell types and other genes to assess its extent, as this may be a broader phenomenon than anticipated. The examples of *Eya3* and *App* demonstrate that some genes can be strictly biallelic. However, *App* was found to be RME in a different cell type^6^, reinforcing the idea that RAExI is cell-type specific. Biallelic genes in one cell type could show RAExI in another. Reciprocally, genes showing RAExI in NPCs could be biallelically expressed in a different context. Therefore, it appears particularly relevant in future studies to assess whether genes associated with autosomal dominant disorders have a RAExI pattern in specific tissues, because it could play a role in the etiology of the disease in a non-canonical manner.

RAExI can be a means to regulate the differential allelic abundance of transcripts, which contributes to the control of gene expression levels in addition to other effects, such as genetic variation. Indeed, genetic polymorphisms at specific loci partly explain variations of gene expression levels (expression quantitative trait loci, eQTLs)^32^. Parent-specific single nucleotide polymorphisms (SNPs) within cis-eQTLs can account for the differential allelic abundance of transcripts, causing a permanent imbalance in expression^33^. RAExI then allows a variability of the allelic imbalance, in a given genetic context. In this study, we have used a 129/Sv x Cast hybrid background *in vitro*, and thus the observed precise modalities reflect the effect of RAExI in this genetic background.

The investigation of RME and RAExI faces technical challenges, as their observation can be confounded with poor detection, genetic aberrations, or transcriptional bursting^14, 17^. In this study, we managed to overcome these limitations. First, we measured the allelic expression ratio in more than 10^6^ cells per clonal cell line, thus averaging out the effect of transcriptional bursts. Moreover, we analysed more than 200 independent NPC clones, which gave us high statistical power and shows that monoallelic expression is not scarce for the genes analysed (it can be present at frequencies >40%). We computed the distribution of the allelic expression ratios, establishing that our observations are not due to technical or biological noise. Furthermore, we ruled out the contribution of genetic aberrations, not only because of the high frequencies of monoallelic expression, but also by showing that the silent allele can be reexpressed for the RME gene *Bag3* using epidrugs. Taken together, our data display robust evidence of RAExI *in vitro*.

*In vivo* analyses are more limited, given the challenge of identifying clonal populations (i.e. cells that originate from the clonal propagation of one cell, after the allelic choice has been established in that cell for the genes of interest). Although there have been some efforts in this direction^6^, *in vivo* analyses are generally performed on single cells and thus, without clonal information. Therefore, these analyses are intrinsically limited by the bursty nature of transcription, which generates transient monoallelic expression or allelic imbalance^31, 32^ that can be confounded with stable RME. Previous studies have observed monoallelic expression *in vivo* by nascent RNA-FISH^11, 19^, but have not excluded the possible contribution of transcriptional bursts. In this study, we developed a RNA-FISH protocol for adult brain, with autofluorescence removal and optimised sensitivity. The implementation of a statistical approach that takes into account the probability of detection, applied to a homogenous cell type in an inbred background, allowed us to show that transcriptional bursts are not responsible for the observed monoallelism of the RME gene *Grik2,* and indicates that stable RME can occur *in vivo*. This has potentially large consequences in physiology and disease^5^. The presence of RME *in vivo* could notably generate some functionally nullisomic cells in a heterozygous context with a loss-of-function mutation. This, for example, is the case of the *Cubulin* gene, which shows mosaicism in the kidney^34^, but it could potentially concern many other genes.

In this study, we also investigated the physiological consequences of allelic expression modalities. We show that these expression patterns generate a wide diversity of gene expression, as all the clones have a unique combination of allelic expression ratios. This diversity could confer adaptability to the organism to environmental perturbations (during development and through life), as some cells could have an advantageous combination of allelic modalities^5^. In addition, we show that monoallelic expression impacts gene dosage, not only at the mRNA level, but also the protein level. The regulation of gene dosage during development could have important implications in cell fate decisions and differentiation, notably if a dosage sensitive transcription factor would play a role in differentiation. For example, it has been shown that the stochastic allelic expression changes of the gene *Bcl11b*, which encodes a factor controlling T-cell commitment, influence T-cell fate timing and development^35^. RME or RAExI could also be a mechanism to fine-tune the levels of expression of specific genes in a particular cell type, such as *Grik2* which shows monoallelism in the CA1 pyramidal neurons, where it is known to be expressed at lower levels^29^. Taken together, we show that RME or RAExI could have some advantageous physiological consequences for an organism.

Despite their potentially major consequences in physiology and pathology, the mechanisms underlying RME are largely unexplored. We show that genes previously categorised as RME do not represent a single homogeneous class of genes, but rather display a wide range of possible allelic states, with some of them showing RAExI rather than RME. The mechanisms of establishment and maintenance of the allelic states are thus most likely gene specific. However, the different genes analysed share common features. First, all genes show biallelic expression in more than half of the NPC clones, indicating that there is no primary feedback selection mechanism to ensure strict monoallelism, thus the expression of both alleles appears to be independent (in contrast to immunoglobulins or olfactory receptors^36^). Monoallelic expression might arise through stochastic silencing or activation of one allele and subsequent clonal propagation. The high frequencies of monoallelism (>40%) nevertheless indicate that the establishment of RME is not a fortuitous and rare event. Furthermore, all genes investigated have different allelic expression states, which are consistent across different cell lines and differentiation experiments. Thus, expression states appear to be pre-determined and gene-specific. This contrasts with the proportions of cells in each state, which vary from one differentiation experiment to another. This variation could be a secondary effect, as some cells might acquire a growth advantage depending on the combination of their allelic choices. Thus, the establishment of the allelic states, by its nature, is stochastic, and has an unpredictable outcome, among the numerous pre-determined possibilities.

Our study addresses more specifically the mechanisms of maintenance of RME. In line with previous observations^13^, we find that the regulation of RME takes place locally at the TSS. Several studies have associated RME with the presence of H3K4me3 and H3K9me3 marks respectively on the active and inactive alleles^10, 13, 37, 38^, not only at the TSS but also on the gene body. Here, we show for the RME genes *Acyp2* and *Bag3*, that the differentially accessible region at the TSS coincides with a CpG island, which is initially unmethylated in ESCs, and is methylated only when the allele is silenced or lowly expressed after differentiation to NPCs. Furthermore, we demonstrate the causality between DNA methylation and the maintenance of monoallelic expression of *Bag3*, as an inhibition of DNA methyltransferases by Decitabine contributes to the derepression of the silent allele. Our epidrug screen also uncovers the role of histone deacetylation in the maintenance of silencing. Hence, using the *Bag3* gene as a proof of principle, we showed that epigenetic modifications contribute to the maintenance of RME. As allelic expression modalities are gene-specific, RME and RAExI genes are likely to depend on specific mechanisms and combinations of epigenetic modifications. The present work should provide a useful set of tools for the future study of other RME and RAExI genes.

We previously found that genes classified as RME are often associated with autosomal dominant disorders in humans and haploinsufficiency in mice^11^. As we observed that the proportions of cells in each allelic state are highly variable for each differentiation experiment, this could contribute to the incomplete penetrance of the disease associated with these genes between individuals. *Bag3,* for example, is associated with autosomal dominant dilated cardiomyopathy with partial penetrance, and has been described as haploinsufficient^39, 40^. We showed that *Bag3* monoallelic expression is maintained by a combination of epigenetic modifications found mostly at the promoter region of the gene. In the case of a loss-of-function heterozygous mutation on a gene expressed in a random monoallelic manner, reactivating the wild-type allele using epidrugs or epigenome editing tools could offer a novel therapeutic strategy for some autosomal dominant disorders. Alternatively, the mechanisms of monoallelic expression could be used to selectively silence a dominant negative allele.

In conclusion, we have uncovered unexpected, various and complex modalities of allelic expression, which go beyond the concept of RME. These modalities of expression may have broad implications in physiology and disease, and represent novel therapeutic perspectives. However, given that each gene appears to show a specific set of expression modalities, it implies that genes should be considered individually rather than by class. We hope that the present work will raise awareness about the concept of epigenetically regulated allele-specific expression, and that it will be considered and investigated in the study of specific genes, for their role both in physiology and pathology, and despite technical challenges.

## METHODS

### Cell culture

The female F1-21.6 and male F1-23 mouse ESC lines were a kind gift from Prof. Joost Gribnau. Their genetic background is hybrid between Mus musculus (129/Sv) and Mus musculus Castaneus (Cast/EiJ). ESCs were grown on mitomycin C-inactivated MEFs in Dubelcco’s Modified Eagle Medium (DMEM) supplemented with fetal bovine serum 15%, β-mercaptoethanol 0.1 mM and leukaemia inhibitory factor 1000 U/mL, at 37 °C with 8% CO2. ESCs were differentiated into NPCs as previously described^11, 21^. Briefly, for each experiment, 1×10^6^ ESCs were plated in a petri dish coated with 0.1% gelatin, in N2B27 medium. After 7 days, cells were dissociated and 3×10^6^ cells were plated in a bacterial petri dish in N2B27 medium supplemented with EGF 10 ng/mL (Peprotech) and FGF2 10 ng/mL (R&D systems), for 3 days. Neurospheres were then transferred in a gelatin-coated petri dish, in the same medium, for them to attach and the NPCs to expand. NPC clones were generated following manual colony picking after limited dilution of the NPC population. Clones were individually expanded and then harvested for RNA, DNA or protein extraction. NPC clones were differentiated in astrocytes as previously described^11^.

### RNA extraction, reverse transcription, qPCR and pyrosequencing

Total RNAs were extracted from NPC cell pellets using the Qiagen RNeasy plus mini kit and treated with DNase1 using the RNase-free DNase set (Qiagen), following the manufacturer’s instructions. Total RNAs (1 µg) were reverse-transcribed into cDNAs using random primers and SuperScript III reverse transcriptase (Invitrogen) for 1 h at 50 °C. Quantitative PCR (qPCR) was performed using Power SybrGreen PCR master mix on a ViiA 7 real-time PCR system (Applied Biosystems). Expression of genes was normalised to three housekeeping genes (*Gapdh*, *B2m*, *Rrrm2*). Clones were classified into “low expression” and “high expression” using an arbitrary threshold (indicated in Figure 5). For pyrosequencing, PCR and sequencing primers were designed using the PyroMark Assay Design software (Qiagen). All successfully amplified PCR products were analysed on a PyroMark Q24 (Qiagen). All primer sequences can be found in Supplementary Tables 7 and 8.

### Definition and classification of the allelic ratios

The allelic expression (for RNA-seq) or accessibility (for ATAC-seq) ratio was defined as the percentage of reads from the Castaneus allele. For sequencing data: (number of reads on the Cast allele) / (number of reads on the Cast allele + number of reads on the 129 allele). For pyrosequencing data: (abundance of sequences from the Cast allele) / (abundance of sequences from the Cast allele + abundance of sequences from the 129 allele). The ratios range from 0% to 100%, corresponding respectively to exclusive monoallelic expression/accessibility from the maternal (129/Sv) allele or the paternal (Cast) allele for a given gene. The allelic categories (for Figure 4b and Figure 5a) are defined as follows: monoallelic [ratio<15% or >85%], biased [ratio 15%-35% or 65%-85%] and biallelic [35%-65%].

### Analysis of allelic expression ratio distributions

The Gaussian mixture models were computed using the mclust package (version 5.4.7)^41^ of the R program, with the EM standard algorithm. The number of components was estimated using the Bayesian Information Criterion (BIC, Supplementary Figure 2). To ascertain that the sample size was sufficient to identify population mixtures with distinct allelic ratios, we further conducted likelihood-ratio tests (LRT) using a bootstrap approach (with 1,000 replications) to evaluate the null distribution and compute p-values^42^ (Supplementary Table 3), using the mclustBootstrapLRT function of the mclust package. We further used a bootstrap approach (1,000 replications again) to calculate 95% confidence intervals for the parameters of the mixture model (Supplementary Table 4) using the MclustBootstrap function^43^.

### Heatmap

The heatmap was performed using XLSTAT (Addinsoft).

### ATAC-seq and RNA-seq

All ATAC-seq datasets (from 13 NPC clones) used for this study were from Xu et al., 2017. Coordinates were lifted over from mm9 to mm10. RNA-seq datasets from 9 NPC clones used in this study were obtained from previously published work^11, 31^. For the additional 7 NPC clones and 5 astrocytes samples, total RNA (1 µg) were prepared and used for polyA enrichment or ribosomal RNA depletion (RiboMinus Eukaryote System V2, Ambion), library preparation (TruSeq Stranded RNA-seq Library Prep, Illumina) and 75 bp or 100 bp paired-end sequencing using a HiSeq2500. All biological replicates were merged. Strain-specific variants (CAST/EiJ and 129S1/SvImJ) were downloaded from the Sanger Mouse Genomes Project (ftp://ftp-mouse.sanger.ac.uk/REL-1505-SNPs_Indels/mgp.v5.merged.snps_all.dbSNP142.vcf). Strain-specific SNPs were used to generate a FASTA file N-masked at high quality homozygous SNPs differing between the two parental strains with SNPsplit (0.3.2). Reads were aligned to the N-masked genome using Tophat (2.1.0)/Bowtie2 (2-2.2.5) with the following parameters: -- reorder -p 8 -D 70 -R 3 -N 0 -L 20 -i S,1,0.50. Reads spanning the N-masked sites were assigned to the maternal or paternal allele using SNPsplit (0.3.2). Pairs for which both reads are assigned to the same parental allele or for which one read is assigned to one parental allele and the other is unassigned are classified as allele-specific. Reads without SNP information, with conflicting allele assignment, or unexpected alleles at polymorphic sites are flagged as unassigned. Read counts were assigned using allele specific and total BAM files with FeatureCounts (1.5.1) with the following parameters: -C -p -s 2. Raw (FASTQ files) and processed data were deposited in GEO under accession number GSE148348. Details about all clones, names and references can be found in Supplementary Table 5.

### Analysis of the RNA-seq in NPCs and astrocytes

Allelic expression ratios (Cast/total allele-specific reads) were calculated for each gene, in each NPC clone and the corresponding astrocyte population. Values with less than 20 allele-specific reads were excluded. Among the 394 genes previously categorised as RME^11^, 195 could be assessed in at least one NPC-astrocyte pair.

### Analysis of the association between ATAC-seq and RNA-seq allelic ratios

For each gene of interest, allelic accessibility and allelic expression ratios (Cast/total) were calculated for each ATAC-seq and RNA seq peak within +/- 3 Mb of the TSS of the gene considered. Peaks with less than 10 allele-specific reads were excluded from the analysis. Only peaks with more than 3 values and at least 2 clones with an expression bias (<33% or >66%) were included in the regression. For each gene, linear regression was performed between the RNA-seq allelic expression ratio and the allelic accessibility ratio for each ATAC-seq peak.

### DNA methylation analysis

Genomic DNA was extracted from NPC cell pellets using the DNeasy blood and tissue kit (Qiagen) and bisulfite-converted using Epitect Bisulfite Kit (Qiagen). Epityper analysis (Sequenom) for *Acyp2* and *Bag3* was performed as previously described^11^. Primer sequences can be found in Supplementary Table 9.

### Western Blot analysis

NPC cell pellets were resuspended in 100 µL RIPA buffer (Tris-HCl 50 mM pH 8.0-8.5, NaCl 150 mM, Triton X-100 1%, Sodium deoxycholate 0.5%, SDS 0.1%) with protease inhibitor (Roche), and sonicated using a Bioruptor (Diagenode, medium power, 3 x 15 sec). Protein concentration was measured using the Bradford assay (Bio-Rad), and adjusted to 1 µg/µL. LDS sample buffer (Invitrogen) and DTT (250 mM) were added, then samples were heated at 95°C for 5 min. 10 µL of each sample were loaded on a NuPAGE 12% Bis-Tris gel, with a protein ladder (PageRuler 26616, Thermo Scientific), and electrophoresis was performed on an Invitrogen system, with a MOPS SDS buffer (NuPAGE, Invitrogen). The transfer to a nitrocellulose membrane was performed using a Bio-Rad system, in Tris 25 mM, Glycine 192 mM, methanol 20%, for 1h at 100V. The membrane was blocked for 1h in TBS-T 0.1%, BSA 5%. Primary antibodies against BAG3 (rabbit polyclonal, 10599-1-AP, Proteintech) and GAPDH (mouse monoclonal, ab9484, Abcam) were incubated overnight at 1/1,000e. Secondary antibodies anti-mouse conjugated with Alexa Fluor 546 (Invitrogen, A-11030) and anti-rabbit conjugated with Alexa Fluor 488 (Invitrogen, A-11034) at 1/5,000e were incubated for 1h at room temperature. The membrane was then washed and imaged with a ChemiDoc MP Imaging System (Bio-Rad). The level of full-length BAG3 was normalised to the level of GAPDH for each clone analysed.

### Epidrug screen

We used an “epigenetics compound library” from Selleckchem, containing 181 compounds (Supplementary Table 6), dissolved in DMSO or water (172 and 9 compounds respectively). NPCs were plated at a density of 7.5×10^4^ cells/cm^2^ in 96-well plates coated with gelatin 0.1%. Six hours later, compounds were added to each well at a final concentration of 10 µM and 0.5% of solvent (DMSO or water). Cells were collected 48 h later for lysis and reverse transcription was performed using the Cells-to-Ct kit (Applied Biosystems) directly in 96-well plates. The allelic expression ratio of *Bag3* was then measured using pyrosequencing after PCR. The 11 best hits were then tested following the same protocol, in 24-well plates. The dose-response curves were performed in 6-well plates, with various concentrations of Dacinostat and Decitabine and a constant concentration of DMSO (0.5%). The final validation was performed in a 6-well plate format, using new batches of Dacinostat and Decitabine (Selleck Chemicals), dissolved in DMSO; cells were treated for 24 h with 10 µM of drugs, in 0.5% DMSO. For assays in 24-well and 6-well plates, cells were harvested and RNAs extracted using the Qiagen RNeasy plus mini kit, as described earlier.

### Mice

The experiments were conducted with 15-week old female C57Bl/6J mice (Charles River, France). The animals were housed in a 12-hour light–dark cycle, in stable conditions of temperature, with food and water *ad libitum*. Experiments were in accordance with the European Community Council Directive 2010/63/EU and approved by the ethics committee of the *Institut Curie* CEEA-IC#118 and authorized by the *Ministère de l’Education Nationale, de l’Enseignement Supérieur et de la Recherche* (APAFIS#8812-2017020611033784 v2).

### RNA-FISH

RNA-FISH probes were prepared by Nick Translation (Abbott) following the manufacturer’s instructions and with a 4 h enzymatic incubation, using Atto-488 conjugated dUTP (Jena Bioscience), and the BAC clones RP23-399D1 and RP24-310A20 (Chori BACPAC Resources Center) for the *App* and the *Grik2* genes, respectively.

Mice were quickly and deeply anesthetized with pentobarbital (500 mg/kg, i.p.) and transcardially perfused with 40 g/L formaldehyde in phosphate-buffered saline (PBS) for 5 min. Brains were dissected and kept at 4°C for 12 h in a PBS solution containing 15% sucrose and 2 mM VRC (vanadium ribonucleoside complex) before being flash frozen in isopentane (1 min at −30°C) and stored at −80°C. Serial coronal sections (30 µm thick) were made using a cryostat (Leica), mounted onto Superfrost Plus slides, and stored at −80°C.

Sections were fixed for 5 min in 30 g/L formaldehyde in PBS, and washed with PBS, 2 mM VRC, at 4°C. They were permeabilized in 0.5% Triton X-100, PBS 1X, 2 mM VRC for 5 min before being washed with 70% ethanol. To remove autofluorescence, sections were incubated in 0.1% (w/v) Sudan Black B, 70% ethanol, for 10 min at RT, then quickly rinsed in 70% ethanol. Sections were equilibrated in 50% (v/v) formamide, 1X SSC (150 mM NaCl, 15 mM sodium citrate), 2.6 mM HCl, pH 7.3, 2 mM VRC, for 30 min at 4°C. 40 ng of labeled DNA probe was precipitated together with 3 µg Cot-1 DNA and 10 µg salmon sperm DNA, before being resuspended in 8 µL formamide, denatured at 75°C, then incubated at 37°C for 30 min for Cot-1 competition. The labeled DNA probe was then hybridized in the presence of Cot-1 and salmon sperm DNA, for 16-24 h at 37°C in 16 µL of the following solution: 50% (v/v) formamide, 1 X SSC, 10 mM VRC, 10% (w/v) dextran sulfate, 0.2% (w/v) BSA. After hybridization, slides were washed two times in 50% formamide/ 2X SSC at 42°C for 30 min, then washed and counterstained with 10 nM TO-PRO-3 in 2X SSC at 42°C for 2 x 30 min. The slides were mounted in Vectashield (Vector Laboratories, USA).

### Microscopy

A laser scanning spinning disk confocal microscope (Roper/Nikon) with a 40× objective was used for image acquisition, which was carried out at the Cell and Tissue Imaging Platform (PICT-IBiSA) at *Institut Curie*.

### Statistical analysis for RNA-FISH

We tested whether the monoallelism measured by RNA-FISH could be explained by transcriptional bursts and/or the sensitivity of detection. We consider that the probability to detect (P_detect_) the nascent RNA for an active allele is:

P_detect_ = P_ON_ x P_sense_, where P_ON_ is the probability to be in a ON phase of transcription, and Psense is the sensitivity of the RNA-FISH, (P_sense_ depends on the power of detection of the RNA-FISH and on the expression level).

For a biallelic gene with identical and unsynchronised alleles, we estimate P_detect_ = n_tot_ / (2 x N_cells_), where n_tot_ is the total number of pinpoints detected over N cells (and 2 alleles per cell).

We reasoned that if the observed monoallelism was due to transcriptional bursts or to limitations of our detection method, the probabilities of detecting each allele should be independent and identical, such that the number of alleles observed would follow a binomial distribution B(2,P_detect_). We tested this hypothesis using a *χ*2 goodness of fit test with one degree of freedom. We reject the binomial hypothesis for *Grik2* in CA1 neurons, which implies that the monoallelism observed for that gene cannot be explained by technical limitations.

### Contributions

AVG, EH and LMP conceptualized the study, designed the experiments and co-supervised this work. AVG, BF, LMP and MA performed *in vitro* experiments. LMP carried out RNA-FISH experiments. BF performed the qPCR and pyrosequencing experiments. AC, AVG, HYC and MA generated RNA-seq data. DZ, FM, LS and LMP analyzed data. AVG, EH and LMP interpreted and discussed the data. AVG, EH and LMP wrote the manuscript, with critical revision of the article by AC, DZ, and HYC. All the authors approved the final version of the manuscript.

## Acknowledgements

We thank Isabelle Grandjean, Colin Jouhanneau and Cédrick Pauchard (In Vivo Experiments platform of *Institut Curie*) for the animal care, Aude Muzerelle for histology advice, Agnès Le Saux and Patricia Diabangouaya for technical assistance, and members of the Heard lab for stimulating discussions. We are grateful to Joost Gribnau (Oncode Institute, Utrecht, NL) for initially providing the F1-21.6 and F1-23 ESC lines. Microscopy and image analysis were carried out at the Cell and Tissue Imaging Platform of the Genetics and Developmental Biology department (UMR3215/U934) of *Institut Curie*, member of France-Bioimaging (ANR-10-INSB-04). High-throughput sequencing was performed by the ICGex NGS platform of the Institut Curie platform and the Stanford Functional Genomics Facility. The ICGEx NGS platform is supported by grants ANR-10-EQPX-03 (Equipex) and ANR-10-INBS-09-08 (France Génomique Consortium) from the ANR (“Investissements d’Avenir” program), the Canceropole Ile-de-France, and the SiRIC-Curie program (SiRIC grant INCa-DGOS-4654). This work was supported by Biogen, and by grants from the *Fondation Vaincre Alzheimer* (FR-17068) and *France Parkinson* (FP 2017) to EH, and by NIH RM1-HG007735 (to HYC). HYC is an Investigator of the Howard Hughes Medical Institute. LMP was funded by a fellowship from the *Fondation pour la Recherche Médicale* (FRM, SPF20151234950).

## SUPPLEMENTARY INFORMATION

**Supplementary Figure 1.**
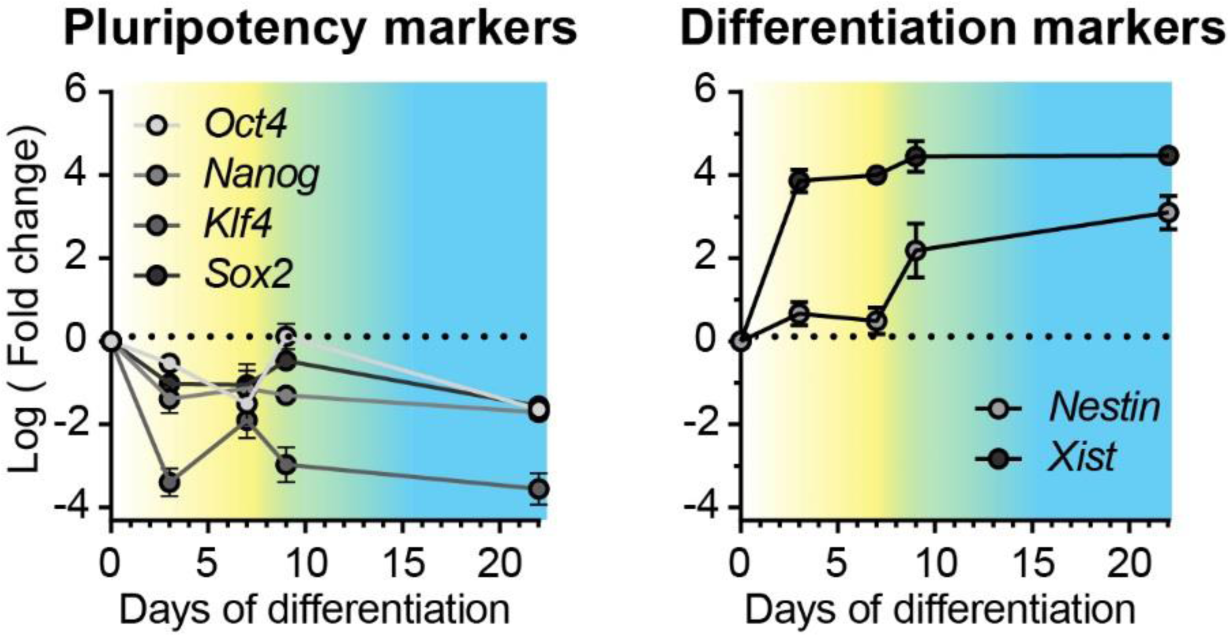
RT-qPCR of pluripotency markers and differentiation markers during the differentiation of ESCs towards NPCs. Data are means ± SEM (*n* = 4 replicates per condition).

**Supplementary Figure 2.**
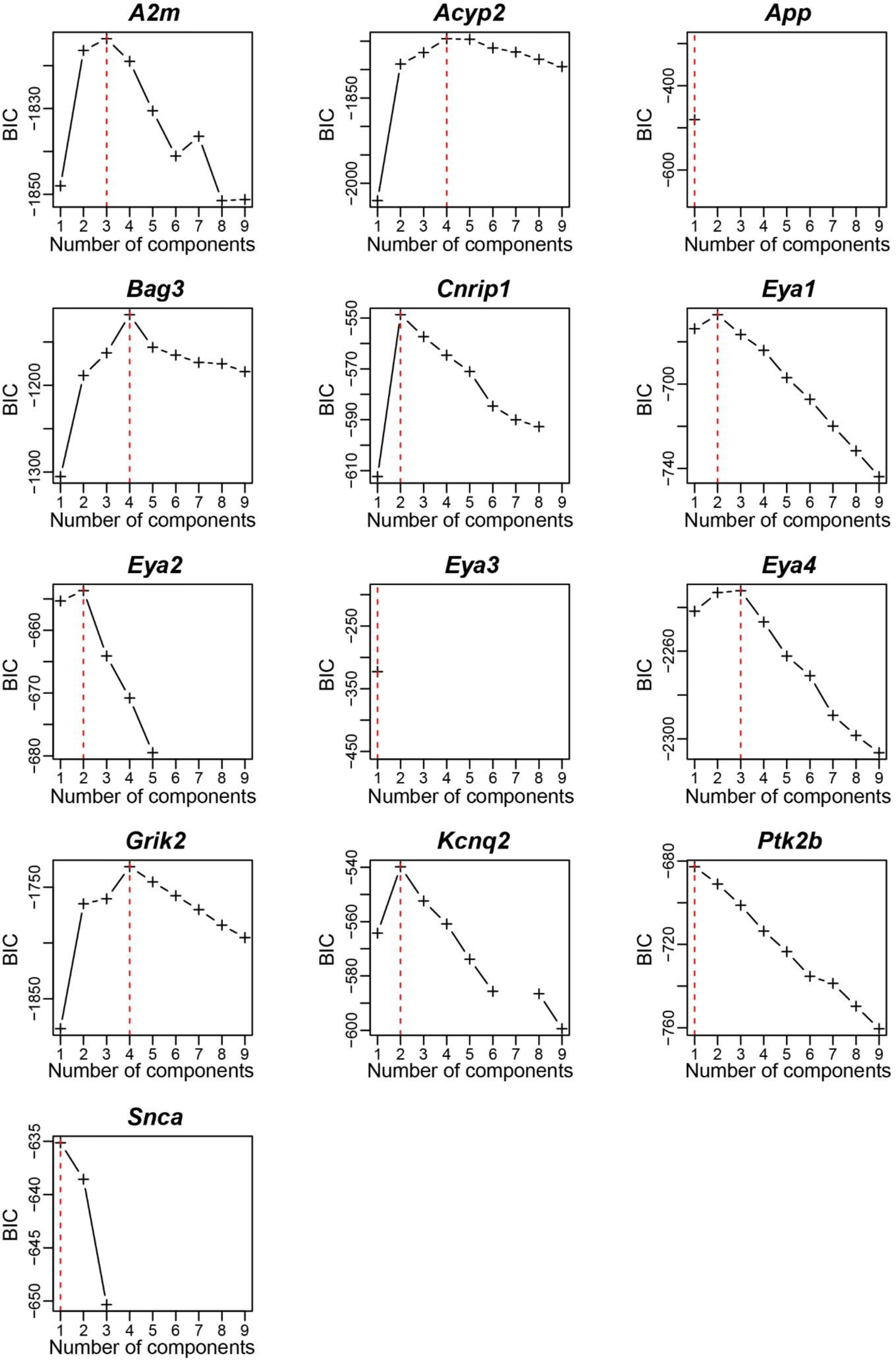
BIC criterion calculated for each gene for the Gaussian mixture models, with up to 9 components. The red dashed lines indicate the number of components maximizing the BIC criterion that was selected. For certain values of the number of components, the EM algorithm did not converge such that the BIC could not be estimated (such as for *Eya3* and *App* - this likely reflects the fact that no good solutions could be found for those number of components).

**Supplementary Figure 3.**
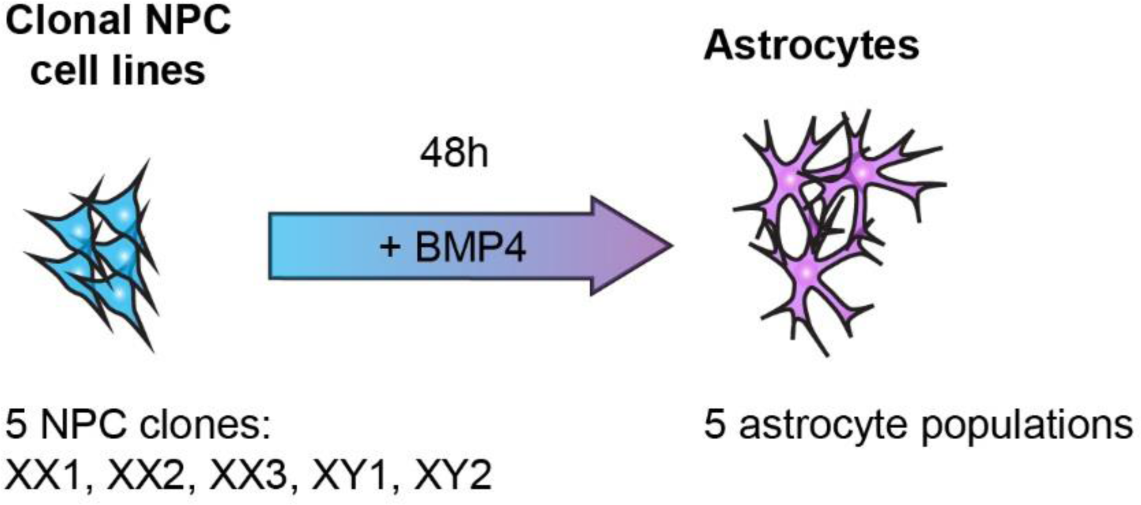
Experimental strategy to assess the stability of monoallelic expression after *in vitro* differentiation of 5 NPC clones into populations of astrocytes.

**Supplementary Figure 4.**
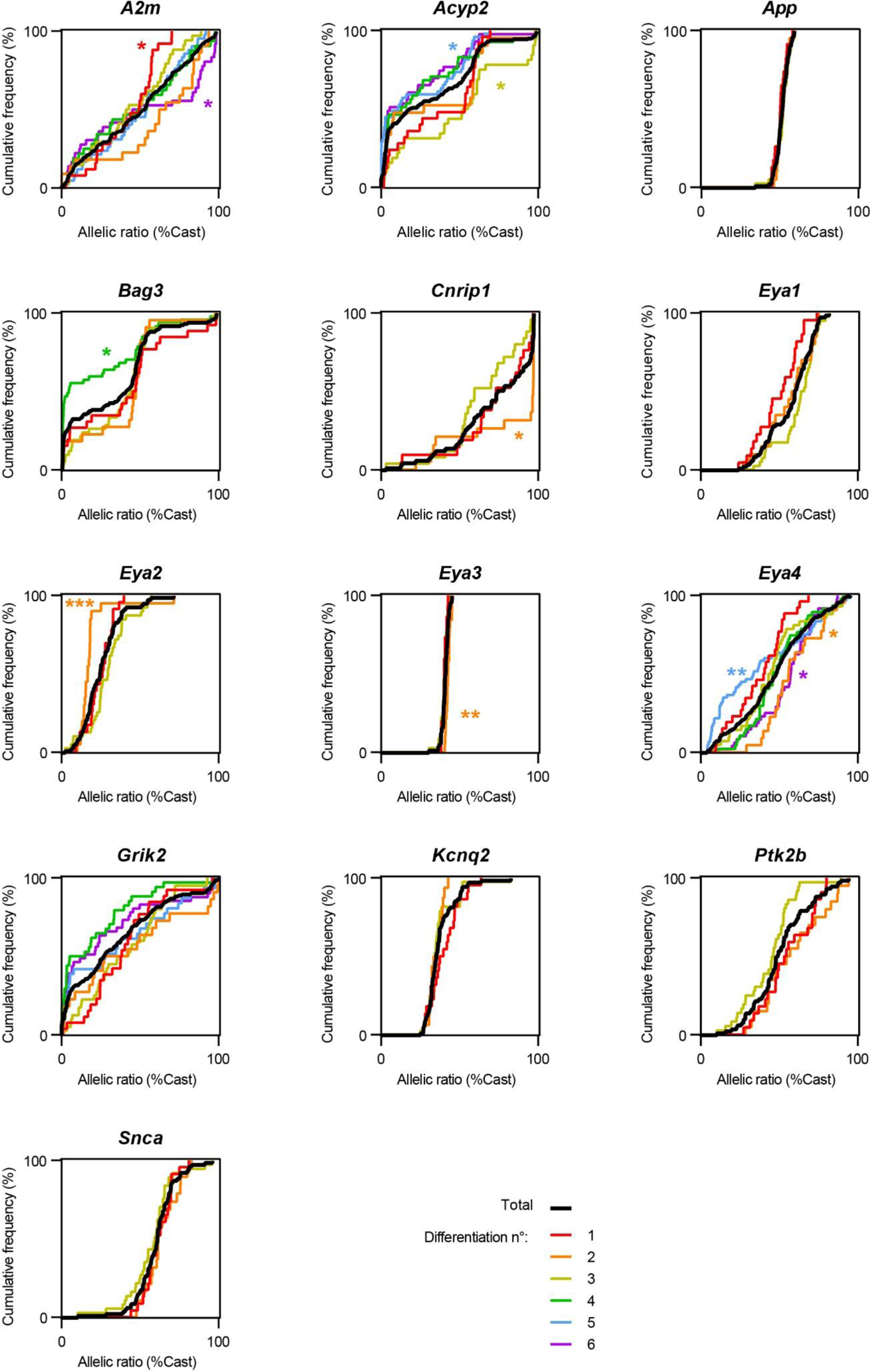
Distributions of allelic expression ratios for each gene of interest, represented as cumulative frequencies, in all NPC clones analysed (in black), and each of the six differentiation experiments independently (colored lines). Kolmogorov-Smirnov tests between the total distribution and the distribution of each experiment, without correction: *p<0.05, **p<0.01, ***p<0.001.

**Supplementary Figure 5.**
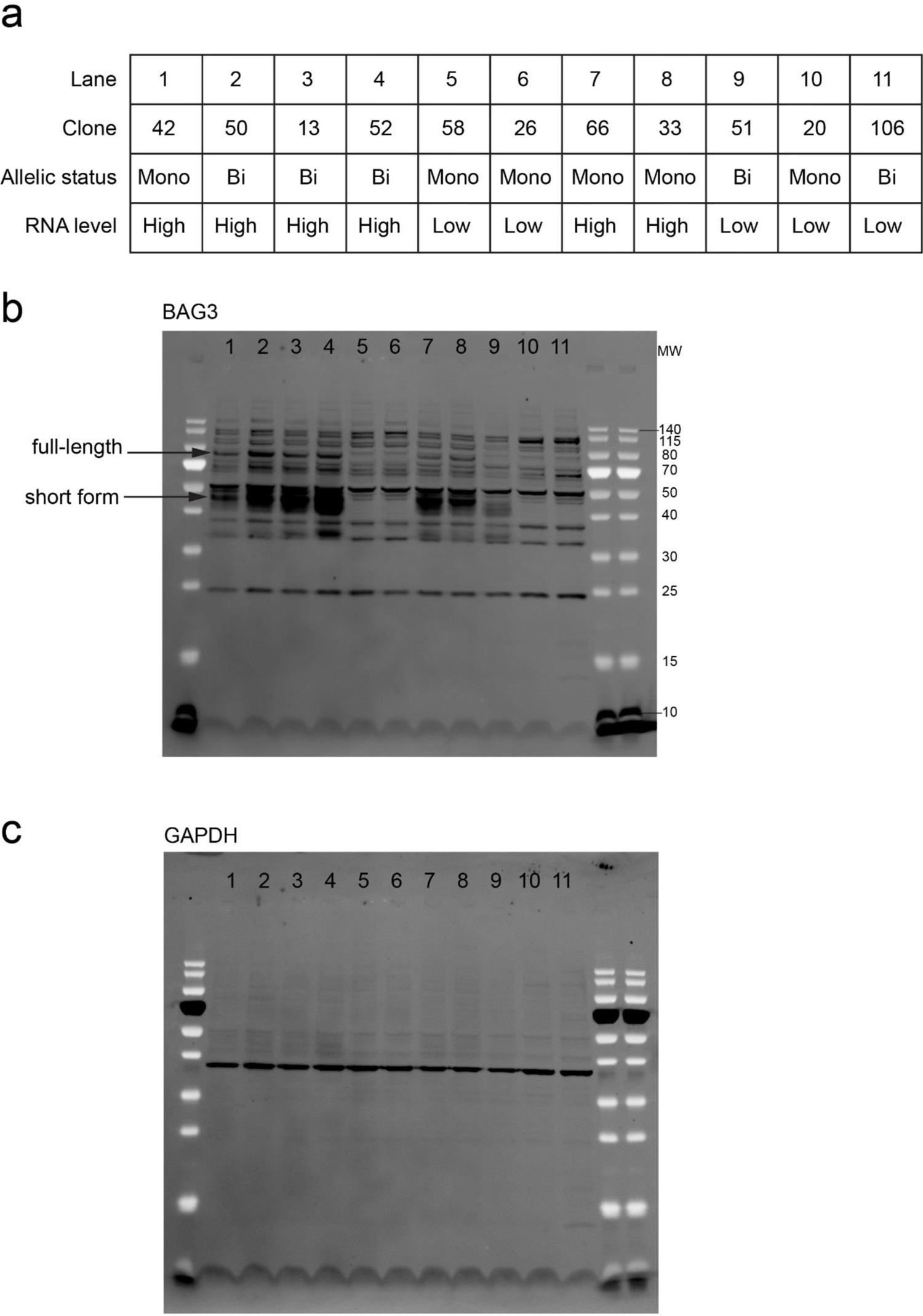
Full image of the western blot analysis of BAG3 protein expression levels. (a) Details of the 11 NPC clones analysed, characterized by high or low expression levels and mono- or biallelic expression. (b) Full image of the western blot with the BAG3 antibody. The full-lenght band was used for the quantification in Figure 5. (c) Full image of the western blot with the GAPDH antibody.

**Supplementary Figure 6.**
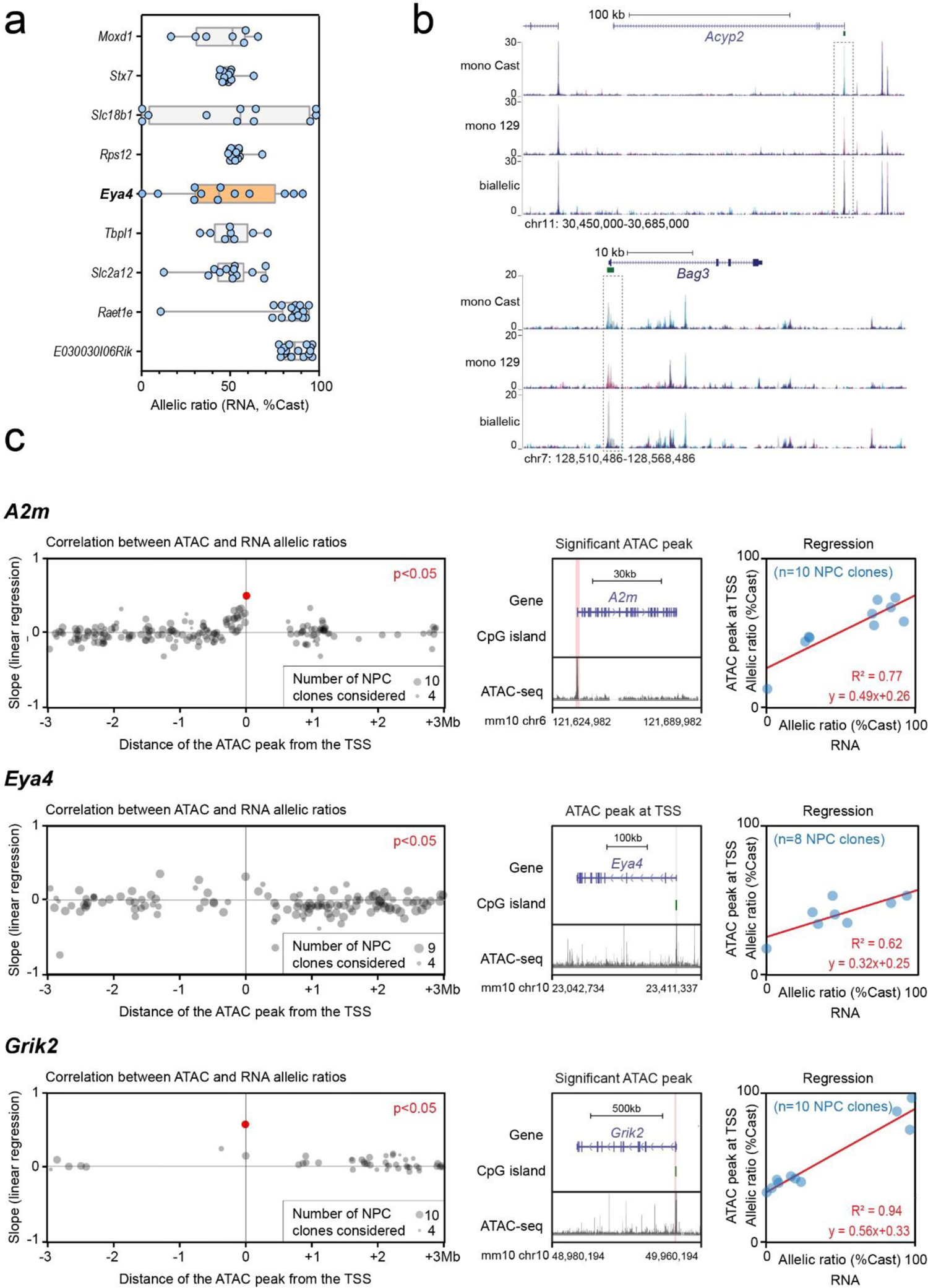
(a) Comparison of the distribution of allelic expression ratios for *Eya4*, with 4 upstream and 4 downstream neighbouring genes, measured by allele-specific RNA-seq in 16 NPC clones. The orange color highlights the whisker plot of the RME gene of interest. (b) Allele-specific ATAC-seq tracks for *Acyp2* and *Bag3* are shown for 3 representative clones (Cast-specific reads are shown in blue, 129-specific reads in pink, and non-allele specific reads in grey). Dashed grey boxes indicate the position of the random accessible monoallelic element for both genes. (c) Identification of the genomic region where allele-specific accessibility correlates best with expression for *A2m*, *Eya4* and *Grik2* by linear regression between the ATAC-seq allelic accessibility ratio and RNA-seq allelic expression ratio, for ATAC-seq peaks (containing a least a SNP) within a region of ±3 megabases around the TSS. *Pvalue* adjusted with Bonferroni correction, number of ATAC peaks: 176/128/47 *A2m*/*Eya4*/*Grik2* (left panels). Representative ATAC-seq track along the genes, with highlighting of the ATAC-seq peak, located at the TSS, that correlates significantly with the RNA-seq for *A2m* and *Grik2* (middle panels). Linear regression showing the correlation between the ATAC-seq allelic accessibility ratio and the RNA-seq allelic expression ratio at the TSS (right panels). ATAC-seq data and RNA-seq data from 13 NPC clones. *A2m F_(1,8)_* = 27, *p* = 8.5 x 10^-4^; *Eya4* F_(1,6)_ = 9.6, *p* = 0.021*; Grik2* F_(1,8)_ = 120, *p* = 4.3 x 10^-6^

---

Supplementary Table 1: Number of NPC clones generated for each differentiation experiment.

Supplementary Table 2: Pyrosequencing data for the 13 genes analysed in NPC clones from differentiation 1 to 6.

Supplementary Table 3: P-values of the likelihood ratio tests. P-values of the likelihood ratio test for each increment of the number of components. A small p-value indicates that likelihood improvement is large enough and is unlikely explained by noise in the data.

Supplementary Table 4: Bootstrap estimates of the parameters. Table presenting the estimates of the parameters (mean, variance and proportions of clones assigned to each population) for each gene and each component. Values in parenthesis give the 95% confidence intervals estimated via resampling (see Methods).

Supplementary Table 5: List of clones used for sequencing. Supplementary Table 6: List of compounds used in the screen.

Supplementary Table 7, 8 and 9: List of primers (pyrosequencing, qPCR, bisulfite)

